# Causal contributions of the domain-general (Multiple Demand) and the language-selective brain networks to perceptual and semantic challenges in speech comprehension

**DOI:** 10.1101/2022.04.12.487989

**Authors:** Lucy J. MacGregor, Rebecca A. Gilbert, Zuzanna Balewski, Daniel J. Mitchell, Sharon W. Erzinclioglu, Jennifer M. Rodd, John Duncan, Evelina Fedorenko, Matthew H. Davis

## Abstract

Listening to spoken language engages domain-general Multiple Demand (MD, fronto-parietal) regions of the human brain, in addition to domain-selective (fronto-temporal) language regions, particularly when comprehension is challenging. However, there is limited evidence that the MD network makes a functional contribution to core aspects of comprehension. In a behavioural study of volunteers (n=19) with chronic brain lesions, but without aphasia, we assessed the causal role of these networks in perceiving, comprehending and adapting to challenging spoken sentences. A first task measured word report for acoustically degraded (noise-vocoded) sentences before and after training. Participants with greater damage to MD but not language regions required more vocoder channels to achieve 50% word report indicating impaired perception. Perception improved following training, reflecting adaptation to acoustic degradation, but perceptual learning was unrelated to lesion location or extent. A second task used sentence coherence judgements to measure the speed and accuracy of comprehension of spoken sentences using lower-frequency meanings of semantically ambiguous words. Comprehension accuracy was high and unaffected by lesion location or extent. The availability of the lower-frequency meaning, as measured in a subsequent word association task, increased following comprehension (word-meaning priming). Word-meaning priming was reduced for participants with greater damage to language but not MD regions. We conclude that language and MD networks make dissociable contributions to challenging speech comprehension: using recent experience to update word meaning preferences depends on language specialised regions, whereas the domain-general MD network plays a causal role in reporting words from degraded speech.

## 2. Introduction

During speech comprehension, listeners are continually challenged by various aspects of the input, which leads to uncertainty at multiple levels of the linguistic hierarchy. For example, acoustic challenges arise when speech is quiet, accented, produced by a young child who has not yet mastered articulation, or otherwise degraded. In such cases, perception of the individual phonemes and lexical forms is more uncertain. Linguistic challenges arise when there is lexical-semantic or syntactic ambiguity, or complexity from low-frequency words or constructions, such that the intended meaning is unclear. To resolve these uncertainties during speech comprehension, listeners make use of diverse sources of information (Altmann & Kamide, 1999; Cutler, Dahan, & van Donselaar, 1997; Garrod & Pickering, 2004; Hagoort, Hald, Bastiaansen, & Petersson, 2004; Münster & Knoeferle, 2018; Özyürek, 2014; Van Berkum, 2009; Zhang, Frassinelli, Tuomainen, Skipper, & Vigliocco, 2021). Furthermore, listeners learn in response to their experiences: they show perceptual and semantic adaptation such that improvements in the perception and comprehension of different types of challenging speech can be observed over time (Davis, Johnsrude, Hervais-Adelman, Taylor, & McGettigan, 2005; Rodd, Cutrin, Kirsch, Millar, & Davis, 2013). In this paper, we consider the potential functional contributions of two distinct groups of cortical brain regions– the domain-selective language network and domain-general Multiple Demand (MD) network - to successful perception and comprehension of different types of challenging speech, and to subsequent perceptual and semantic adaptation.

### 2.1. Role of language-selective versus domain-general (Multiple Demand) regions in language comprehension

The language-selective network is a set of left-lateralised frontal and temporal regions that respond reliably to linguistic stimuli with different input modalities, languages and tasks (e.g., Binder et al., 1997; Blank, Kanwisher, & Fedorenko, 2014; Fedorenko, Duncan, & Kanwisher, 2012; Fedorenko, Hsieh, Nieto-Castanon, Whitfield-Gabrieli, & Kanwisher, 2010; MacSweeney et al., 2002; Mahowald & Fedorenko, 2016; Mineroff, Blank, Mahowald, & Fedorenko, 2018; Paunov, Blank, & Fedorenko, 2019; Scott, Gallee, & Fedorenko, 2017; for a review, see Fedorenko, 2014), but not to non-linguistic stimuli such as music, mathematical expressions, or computer code (Fedorenko, Behr, & Kanwisher, 2011; Ivanova et al., 2020; Monti, Parsons, & Osherson, 2012). These regions are functionally connected (Braga, DiNicola, Becker, & Buckner, 2020), and show correlated response profiles at rest and during naturalistic listening (Blank et al., 2014; Braga et al., 2020; Mineroff et al., 2018; Paunov et al., 2019), leading to their characterisation as a functionally coherent network. Lesion studies show that damage to, or degeneration of, this network leads to impairments in language function (Bates et al., 2003; Mesulam et al., 2014; Mirman et al., 2015; Mirman & Thye, 2018; Turken & Dronkers, 2011), but does not cause deficits in other cognitive domains (Apperly, Samson, Carroll, Hussain, & Humphreys, 2006; Fedorenko & Varley, 2016; Ivanova et al., 2021; Polk & Kertesz, 1993; Varley, Klessinger, Romanowski, & Siegal, 2005; Varley & Siegal, 2000; Varley, Siegal, & Want, 2001) indicating a necessary and selective role of the network in language comprehension.

Sometimes, linguistic stimuli also activate a set of bilateral frontal, parietal, cingular and opercular regions (for a large scale fMRI investigation and relevant discussion, see Diachek, Blank, Siegelman, Affourtit, & Fedorenko, 2020), which together form the Multiple Demand (MD) network (Duncan, 2010b, 2013). This network is domain-general, responding during diverse demanding tasks (Duncan & Owen, 2000; Fedorenko et al., 2012; Fedorenko, Duncan, & Kanwisher, 2013; Hugdahl, Raichle, Mitra, & Specht, 2015; Shashidhara, Mitchell, Erez, & Duncan, 2019) and has been linked to cognitive constructs such as executive control, working memory, selective attention, and fluid intelligence (Assem, Blank, Mineroff, Ademoglu, & Fedorenko, 2020; Cole & Schneider, 2007; Duncan & Owen, 2000; Vincent, Kahn, Snyder, Raichle, & Buckner, 2008; Woolgar, Duncan, Manes, & Fedorenko, 2018). These regions of the MD network show strongly synchronized activity, and fluctuation patterns that dissociate sharply from those of the language network (Blank et al., 2014; Mineroff et al., 2018; Paunov et al., 2019). Moreover, damage to the MD network leads to patterns of cognitive impairment that differ from those observed in cases of language network damage (Duncan, 2010a; Fedorenko & Varley, 2016; Woolgar et al., 2018; Woolgar et al., 2010), confirming a functional dissociation between the two networks (see Fedorenko & Blank, 2020 for a review focusing on the dissociation between subregions of Broca’s area).

Recently, it has been argued that the MD network does not play a functional role in language comprehension (Blank & Fedorenko, 2017; Diachek et al., 2020; Shain, Blank, van Schijndel, Schuler, & Fedorenko, 2020; Wehbe et al., 2021; for reviews, see Campbell & Tyler, 2018; Fedorenko & Shain, 2021). Instead, it is proposed that activation of MD regions reflects a general increase in effort, which is imposed by task demands in particular, or in some cases even mis-localisation of language-selective activity because of the proximity of the two systems in some parts of the brain (e.g., in the inferior frontal gyrus (IFG), Fedorenko et al., 2012; see Quillen, Yen, & Wilson, 2021 for evidence that increased linguistic and non-linguistic task demands matched on difficulty, differentially recruite langage-selective versus domain-general regions).

However, existing evidence that domain-general MD regions do not contribute to language comprehension is limited in two ways. First, relevant studies have typically drawn conclusions about function based on the magnitude of neural activity (e.g., the BOLD fMRI response). The strongest *causal* inference about the necessity (and selectivity) of brain regions for particular cognitive processes comes from approaches that transiently disrupt neural functioning in the healthy brain (e.g., TMS) and measure the effects on behaviour, or from cases of acquired brain damage, either in case studies or multi-patient lesion-symptom mapping investigations that exploit inter-individual variability in behavioural and neural profiles to link specific brain systems to behavioural outcomes (Halai, Woollams, & Lambon Ralph, 2017). A recent lesion study found that the extent of damage to the MD network predicted deficits in fluid intelligence; in contrast, MD lesions did not predict remaining deficits in verbal fluency after the influence of fluid intelligence was removed, which instead were predicted by damage to the language-selective network (Woolgar et al., 2018), in line with the dissociation discussed above. These results provide convincing evidence for a specific contribution of the MD network but not the language network to fluid intelligence. However, given that language function was assessed with a verbal fluency task—an elicited production paradigm that relies on a host of diverse cognitive operations—the question of whether the MD network causally contributes to specific aspects of language comprehension remains unanswered.

A second limitation of previous studies is their focus on the comprehension of clearly perceptible and relatively unambiguous language, whereas naturalistic speech comprehension typically involves dealing with noise and uncertainty in the input. For example, speech may be in an unfamiliar accent, or contain disfluencies and mispronunciations; there may be background speech, other sounds or distractions; and the words and syntax may be ambiguous or uncommon. These features can make identifying words and inferring meaning– core computations of comprehension – more difficult (for a review of different types of challenges to speech comprehension, see Johnsrude & Rodd, 2015). It therefore remains a possibility that the MD network is functionally critical in these more challenging listening situations (Diachek et al., 2020).

### 2.2. Challenges to speech perception and comprehension

Here, we focus on two challenges, which arise from acoustic degradation and from lexical-semantic ambiguity. Acoustically degraded speech makes word recognition less accurate (Mattys, Davis, Bradlow, & Scott, 2012), reduces perceived clarity (Sohoglu, Peelle, Carlyon, & Davis, 2014), increases listening effort (Pichora-Fuller et al., 2016; Wild et al., 2012), and encourages listeners to utilise informative semantic contextual cues (Davis, Ford, Kherif, & Johnsrude, 2011; Miller, Heise, & Lichten, 1951; Obleser, Wise, Dresner, & Scott, 2007; Rysop, Schmitt, Obleser, & Hartwigsen, 2021) and other forms of prior knowledge (Miller & Isard, 1963; Sohoglu et al., 2014; Sumby & Pollack, 1954). Acoustically degraded speech engages brain regions that plausibly fall within the MD network including parts of the premotor, motor, opercular and insular cortex (Davis & Johnsrude, 2003; Du, Buchsbaum, Grady, & Alain, 2014, 2016; Erb, Henry, Eisner, & Obleser, 2013; Evans & Davis, 2015; Hardy et al., 2018; Hervais-Adelman, Carlyon, Johnsrude, & Davis, 2012; Vaden, Kuchinsky, Ahlstrom, Dubno, & Eckert, 2015; Vaden et al., 2013; Wild et al., 2012), as well in the angular gyrus (Rysop et al., 2021) and inferior frontal gyrus (Davis et al., 2011; Davis & Johnsrude, 2003). Furthermore, disruption of premotor regions either by TMS (D’Ausilio et al., 2009; D’Ausilio, Bufalari, Salmas, & Fadiga, 2012; Meister, Wilson, Deblieck, Wu, & Iacoboni, 2007) or following lesions (Moineau, Dronkers, & Bates, 2005; for a review, see Pulvermuller & Fadiga, 2010) has been shown to impair perception of degraded speech. However, given that the MD network was not explicitly defined in previous studies, the functional contribution of the MD network to acoustically degraded speech perception remains untested.

Lexical-semantic ambiguity (for a review, see Rodd, 2018) challenges comprehension because of the competition between alternative meanings of a single word form during meaning access (Rayner & Duffy, 1986; Rodd, Gaskell, & Marslen-Wilson, 2002; Seidenberg, Tanenhaus, Leiman, & Bienkowski, 1982; Swinney, 1979), and because costly reinterpretation is sometimes required (Blott, Rodd, Ferreira, & Warren, 2021; Duffy, Morris, & Rayner, 1988; Rodd, Johnsrude, & Davis, 2010, 2012). Domain-general cognitive operations may be useful in responding to the challenge, as evidenced by the positive relationship between an individual’s success in semantic ambiguity resolution and executive functioning skill (Gernsbacher & Faust, 1991; Gernsbacher, Varner, & Faust, 1990; Khanna & Boland, 2010) and dual task studies showing that performance on non-linguistic visual tasks is impaired during semantic reinterpretation (Rodd et al., 2010), but these domain-general operations may be plausibly generated by either language-selective or domain-general cortical regions.

Functional imaging studies show that semantic ambiguity resolution engages left-lateralised frontal and temporal brain regions typical of the language-selective network, specifically posterior parts of middle and inferior temporal lobe, anterior temporal lobe, and the posterior IFG (Bilenko, Grindrod, Myers, & Blumstein, 2009; Musz & Thompson-Schill, 2017; Rodd, Davis, & Johnsrude, 2005; Vitello, Warren, Devlin, & Rodd, 2014; Zempleni, Renken, Hoeks, Hoogduin, & Stowe, 2007; for a review, see Rodd, 2020). The possibility that the IFG in particular plays a causal role is supported by the observation that individuals with Broca’s aphasia have difficulties in using context to access subordinate word meanings (Hagoort, 1993; Swaab, Brown, & Hagoort, 1997; Swinney, Zurif, & Nicol, 1989), although patients in these studies were selected based on their language profile rather than lesion location.

Although subregions within the IFG form part of the language-selective network, as discussed above, there are also subregions that fall within the domain-general MD network (e.g., Fedorenko & Blank, 2020). Indeed IFG recruitment during ambiguity resolution has been typically accounted for by invoking domain-general constructs of cognitive control or conflict resolution (Novick, Trueswell, & Thompson-Schill, 2005; Thompson-Schill, D’Esposito, Aguirre, & Farah, 1997) which resolve competition between alternative meanings of ambiguous words (Musz & Thompson-Schill, 2017). Currently, the heterogeneity of the IFG makes activations within this region difficult to interpret functionally, without careful anatomical identification of relevant components (Tahmasebi et al., 2012).

A range of studies show that listeners can adapt to the challenges of perceiving and comprehending acoustically degraded or semantically ambiguous sentences. Listeners’ perception of degraded speech improves spontaneously over time with repeated exposure (Davis et al., 2005; Guediche, Blumstein, Fiez, & Holt, 2014; Hervais-Adelman, Davis, Johnsrude, & Carlyon, 2008; Loebach & Pisoni, 2008; Sohoglu & Davis, 2016; Stacey & Summerfield, 2008), so long as attention is directed to speech (Huyck & Johnsrude, 2012). This perceptual adaptation is facilitated by visual/auditory feedback presented concurrently or in advance (Wild et al., 2012), generalises across talkers (Huyck, Smith, Hawkins, & Johnsrude, 2017), is supported by lexical-level information such that learning through exposure to pseudo-words is less effective than with real words (although in some cases, learning with pseudowords is possible, Hervais-Adelman et al., 2008), but does not additionally benefit from sentence-level semantic information (learning was as effective with meaningless syntactic prose; Davis et al., 2005).

Regarding adaptation to ambiguous words, research has shown that accessing a less frequent (subordinate) meaning of an ambiguous word is easier following exposure to the same meaning of an ambiguous word in a prime sentence. Although the cognitive operations underpinning this so-called word meaning priming effect remain somewhat underspecified, the effect can be described as a form of longer-term lexico-semantic learning since it can be observed tens of minutes or even hours after initial exposure, or perhaps longer if adaptation is consolidated by sleep (Betts, Gilbert, Cai, Okedara, & Rodd, 2018; Gaskell, Cairney, & Rodd, 2019; Rodd et al., 2013).

### 2.3. The current study

In the current study, we ask whether speech perception and comprehension in these different challenging circumstances, and adaptation in response to these challenges, depend on the MD network or the language-selective network. To do this, we investigated the impact of lesions to these networks, on word recognition and interpretation – two core components of language comprehension – and on adaptation, using behavioural measures.

Participants (n=19) with long-standing lesions to the domain-selective language network and/or the domain-general Multiple Demand (MD) network performed two behavioural tasks, each of which had two phases to assess both immediate effects and longer-term consequences of the specific listening challenge. Task 1 (acoustic-phonetic challenge) measured perception of noise-vocoded spoken sentences in a word report task and the consequences of training with this type of acoustic degradation for subsequent perception. Task 2 (lexico-semantic challenge) measured comprehension of spoken sentences that included semantically ambiguous words in a sentence coherence judgement task. The consequences of experience with lower-frequency meanings of ambiguous words for subsequent meaning access were assessed in a word association task.

For each task, behavioural performance measures were associated with lesion location and extent by performing correlational analyses using probabilistic functional activation atlases (e.g., Woolgar et al., 2018). Finally, across-task analyses assessed potential dissociations between the contributions of these two networks for accommodating and adapting to different sources of listening challenge during speech comprehension.

## 3. Methods

### 3.1. Participants

Twenty-one right-handed native English speakers were recruited from [DETAIL REMOVED FOR DOUBLE BLIND REVIEWING], a database of volunteers who have suffered a brain lesion and have expressed interest in taking part in research. Participants were invited to take part in the current research on the basis that they had chronic lesions (minimum time since injury of 3 years) to cortical areas falling predominantly in the language or Multiple Demand (MD) networks (or lesions in other areas for control participants). These networks were broadly defined based on previous functional imaging data from typical volunteers (described below), and linked to lesions traced on anatomical MRI scans for [DETAIL REMOVED FOR DOUBLE BLIND REVIEWING] volunteers selected without knowledge of their behavioural profiles. Thus, volunteers were not recruited on the basis of a known language impairment or aphasia diagnosis. Participants gave written informed consent under the approval of [DETAIL REMOVED FOR DOUBLE BLIND REVIEWING]. Data from two participants were not included in the final analyses of either task (one participant was unable to complete either task due to fatigue and hearing difficulties; a second failed to complete the semantic ambiguity experimental tasks and also had difficulties accurately reporting back the words they heard in the degraded speech task, achieving only 68% word report accuracy for the clear speech condition across pre- and post-training test sessions, see task details below).

The remaining 19 participants (8 female, mean age 61 years, range 37 – 75 years) had brain lesions caused by tumour excision (n=8), stroke (haemorrhagic: n=6, ischaemic: n=1), or a combination of these (tumour excision and haemorrhagic stroke: n=1) with other causes being abscess excision (n=1) or resection because of epileptic seizures (n=1), and one of unknown cause (n=1). Individual participant characteristics are detailed in Table 1. Two participants contributed data to just one of the tasks (md6 was excluded from the degraded speech experiment for not completing the task; md10 was excluded from the semantic ambiguity analyses for giving multiple responses during the word association task). Thus, data from 18 participants were included for each of the experiments analysed separately (see below) and from 17 participants for the cross-experimental analyses. The National Adult Reading Test (NART) (Nelson, 1982) was used to estimate premorbid IQ. The Test of Reception of Grammar (TROG-2, Bishop, 2003) was used as a background assessment of linguistic competence.

**Table 1.** Participant profiles including demographics, lesion details, IQ based on the NART and TROG2 scores. Participant md6 was excluded from the degraded speech task analyses; participant md10 was excluded from the semantic ambiguity task analyses.

| Participant | Group | age at<br>test | sex | Lesion<br>aetiology | Lesion<br>location | Premorbid<br>IQ (from<br>NART) | TROG2<br>Score | Total<br>Lesion<br>Volume<br>(cm<sup>3</sup>) |
| --- | --- | --- | --- | --- | --- | --- | --- | --- |
| lang1 | LANG | 50 | F | Tumour | L posterior<br>temporal,<br>posterior corpus<br>callosum,<br>posterior<br>cingulum | 108 | 14 | 29.49 |
| 88lang2 | LANG | 75 | F | Stroke (ischaemic) | L frontal and anterior insular<br>cortex | 86 | 8 | 64.06 |
| lang3 | LANG | 58 | F | Stroke<br>(haemorrhagic) | R fronto-temporo-<br>parietal and<br>anterior thalamus | 123 | 18 | 139.70 |
| lang4 | LANG | 65 | F | Tumour | L temporal | 117 | 16 | 21.90 |
| lang5 | LANG | 50 | M | Other (abscess<br>removal) | L temporal | 101 | 14 | 19.71 |
| lang6 | LANG | 56 | M | Tumour | R inferior<br>parietal/temporal | 112 | 12 | 42.38 |
| md1 | MD | 58 | M | Stroke<br>(ischaemic) | L occipito-parietal | 97 | 15 | 78.21 |
| md2 | MD | 63 | F | Tumour | L superior parietal<br>lobe | 123 | 17 | 37.06 |
| md3 | MD | 64 | M | Other<br>(unknown) | R fronto-temporo-<br>parietal<br>surrounding<br>insular cortex.<br>Anterior branch of<br>internal capsule | 115 | 15 | 68.59 |
| md4 | MD | 53 | F | Tumour | R superior frontal | 106 | 14 | 114.68 |
| md5 | MD | 69 | M | Stroke<br>(haemorrhagic) | L frontal | 101 | 4 | 113.15 |
| md6* | MD | 61 | M | Tumour +<br>Stroke<br>(haemorrhagic) | R posterior frontal<br>and some<br>anterior/medial,<br>extending to post-<br>central<br>gyrus/parietal<br>areas | 123 | 13 | 131.27 |
| md7 | MD | 52 | F | Tumour | Bifrontal | 112 | 16 | 59.02 |
| md8 | MD | 74 | M | Tumour | R frontal | 120 | 16 | 27.68 |
| md9 | MD | 67 | M | Tumour | L anterior frontal | 126 | 18 | 24.05 |
| md10* | MD | 81 | M | Stroke<br>(haemorrhagic) | L temporo-<br>occipital | 81 | 14 | 13.26 |
| other1 | OTHER | 58 | M | Other (resection<br>for epilepsy) | R temporal | NA | 16 | 17.84 |
| other2 | OTHER | 60 | M | Stroke<br>(haemorrhagic) | R basal ganglia<br>(putamen +<br>caudate +<br>thalamus) and<br>internal capsule<br>dorsal anterior<br>insula | 97 | 17 | 25.37 |
| other3 | OTHER1 | 37 | F | Stroke<br>(haemorrhagic) | L frontal | 105 | 15 | 12.52 |

### 3.2. Lesion analysis

Lesion analysis followed procedures developed in previous research and here we only report key points (for further details see Woolgar et al., 2018). Each participant had a structural MRI image (T1-weighted Spoiled Gradient Echo (SPGR) MRI scans with 1×1×1mm resolution) which included lesion tracing as part of previous participation in [DETAIL REMOVED FOR DOUBLE BLIND REVIEWING]. From these images, we estimated the volume of lesion that overlapped with the language network, the Multiple Demand network or elsewhere. The two networks were defined from probabilistic fMRI activation maps constructed from large numbers of healthy participants (Language: n=220, MD: n=63) who performed tasks developed to localise language processing and domain-general executive processing (see Blank et al., 2014; Fedorenko, 2014; Fedorenko et al., 2013; Mahowald & Fedorenko, 2016). The activation maps for the language network contrasted data from participants reading or listening to sentences versus lists of pseudowords (neural responses in the language network are modality-independent; Fedorenko et al., 2010; Scott et al., 2017); those for the Multiple Demand network contrasted data from participants performing a hard versus easy spatial working memory task (remembering 8 vs. 4 locations, respectively, in a 3 × 4 grid). Each individual participant’s activation map for the relevant contrast (sentences > pseudowords, hard > easy spatial working memory) was thresholded at a p< .001 uncorrected level, binarised and normalised before the resulting images were combined in template space. The resulting probabilistic activation overlap maps (Figure 1A) contain information in each voxel about the proportion of participants who show a significant effect (at p< .001) for the contrast of interest. Following Woolgar et al. (2018), voxels in which fewer than 5% of the contributing participants showed a significant effect were excluded.

**Figure 1.**
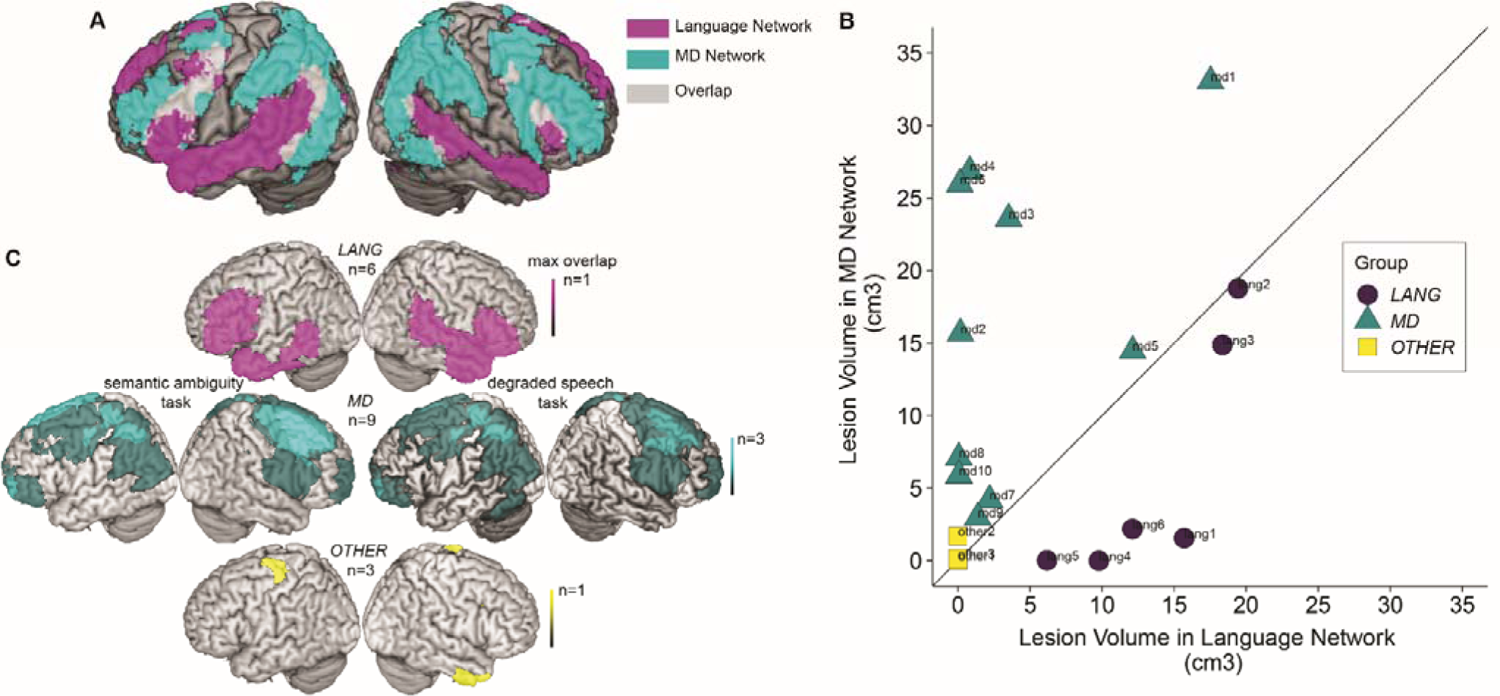
A.The language and MD networks against which we compared participants’ lesions. The images show probabilistic activation maps of the language network and the MD network based on fMRI data from large numbers of neurotypical participants (language: n=220; MD: n=63), which have been thresholded to show regions active in at least 5% of participants during the relevant functional task and plotted onto a volume rendering of the brain. B. Volume of lesion falling into each network for each of the 19 participants in the present study. Solid line depicts an equal volume of each network affected by the lesion. Different colours/shapes indicate assignment of the participants into the *LANGUAGE* (*LANG*), *MD* and OTHER Groups for categorical analyses (see supplementary materials). C. Lesion overlap across participants depicted on volume rendering of the brain. Images are shown separately for participants in the three groups (see supplementary materials) and the two tasks. Brighter colours reflect greater lesion overlap across participants.

We then calculated the lesion volume falling into each network for each of the 19 participants (Figure 1B). Participants were assigned to one of three broad groups (LANG, MD or OTHER) based on the proportion and volume of their lesions falling into language and Multiple Demand regions as well as the overall proportion of each network that was damaged (Figure 1C; see supplementary materials for further details of group assignment). We use these group assignments in describing participants and results; however, since the group allocation for several participants was not clear-cut, we focus on correlational analyses that relate behavioural performance measures and lesion location or volume. Group analyses are included in the supplementary materials.

### 3.3. Statistical analysis

Analyses were performed using R statistical software (version 3.6.1, R Core R Development Core Team, 2019). For each task, the primary analyses assessed whether more extensive damage to the language and MD networks were associated with more impaired performance on the behavioural tasks, with one-tailed Pearson’s r correlation coefficients. We compared the strength of different correlations within task (i.e. comparing the impact of damage to language and MD networks on a given behavioural measure) and between task (i.e. comparing the impact of damage to a given network on different behavioural measures) with two-tailed Meng’s z tests (Meng, Rosenthal, & Rubin, 1992) using the ‘cocor’ package (Diedenhofen & Musch, 2015). The between-task comparisons focused on the 17 participants for whom we had data for both the degraded speech and the lexical-semantic ambiguity tasks.

### 3.4. Task 1. Acoustically degraded speech perception and adaptation

The first task increased speech comprehension difficulty at the acoustic-phonetic level of the input, by acoustic degradation of spoken sentences with noise vocoding (Shannon, Zeng, Kamath, Wygonski, & Ekelid, 1995). Noise vocoding reduces the spectral detail in the speech signal but retains the slow amplitude modulations which approximately reflect syllabic units, and the broad-band spectral changes that convey speech content. These low frequency modulations and broadband spectral modulations have been shown to be most important for accurate speech perception (Elliott & Theunissen, 2009; Shannon et al., 1995). We selected the particular numbers of channels in the vocoder based on previous research, which established that intelligibility (as measured by word report: how many words of the sentence a participant can accurately report) increases with the logarithmic increase in number of channels (McGettigan, Rosen, & Scott, 2014). In healthy adults with good hearing, for short sentences of 6-13 words long, intelligibility is near 100% for 16-channel vocoded speech, near 0% for 1-channel vocoded speech, and at an intermediate level for 4-channel vocoded speech (Peelle, Gross, & Davis, 2013). We assessed speech perception in terms of the logarithmic number of channels estimated as required to achieve 50% word report accuracy of these sentences and assessed adaptation by comparing performance before and after a training period.

#### 3.4.1. Stimuli

The stimuli for the degraded speech task were forty declarative sentences, varying in length (6-13 words, M=9, SD = 2.45) and duration (1.14 to 3.79 seconds, M= 2.12, SD = 0.60), which were selected from coherent low ambiguity sentences used in previous studies (Davis et al., 2011). Sentences were recorded by a female native speaker of British English and digitised at a sampling rate of 22050Hz. We created three degraded versions of the sentences, of decreasing intelligibility, using 16-, 8- and 4-channels in the vocoder. To do this, the frequency range 50-8000Hz was divided into 16, 8, or 4 logarithmically spaced frequency channels. Each channel was low pass filtered at 30Hz and half-wave rectified to produce an amplitude envelope for each channel, which was then applied to white noise that was filtered in the same frequency band. Finally, the channels were recombined to create the noise-vocoded version of the sentence.

The 40 sentences were grouped into 8 sets of 5 sentences such that each set contained 45 words in total and expected (based on previous word report data) to be approximately equally intelligible. Each participant heard all 8 sentence sets, but assignment of sets to the different levels of degradation (clear, 16-, 8-, 4-channel vocoded) and to the pre- and post-training test (described below) was counterbalanced across participants.

#### 3.4.2. Procedure

The experiment started with 4 practice trials to familiarise the participants with the stimuli and the word report task. Participants listened to 4 different sentences (not included in the test set) at decreasing levels of degradation (clear, 16, 8, 4) and after each sentence had to repeat the sentence or as many words from the sentence as possible in the correct order. The experiment then followed a test-train-test format (c.f. Sohoglu & Davis, 2016) with the 40 experimental sentences (8 sets of 5 sentences; see stimuli for details of assignment of the sentences to the pre-test and post-test and to the different levels of degradation). In the initial test, participants listened to 20 of the sentences, 5 at each level of degradation (clear, 16, 8, 4; order randomised uniquely for each participant) and performed the word report task. This was followed by a training period in which participants listened passively to the same 20 sentences, each repeated 4 times at decreasing levels of degradation, whilst the written text of the sentence was presented visually on a computer screen. Following the training, participants listened to the other (previously unheard) 20 sentences, 5 at each level of degradation and again performed the word report task.

#### 3.4.3. Data processing and analysis

For each participant, we calculated the proportion of words correctly reported (total 45 words) at each level of degradation (clear, 16-, 8-, 4-channel vocoded) and for the pre- and post-training test. Words were scored correct only if there was a perfect match with the spoken word from the sentence (morphological variants were scored as incorrect, but homonyms, even if semantically anomalous, were scored correct). Words reported in the correct order were scored correct even if intervening words were absent or incorrectly reported, but scored as incorrect if they were reported in the wrong order. To verify that decreasing the number of vocoded channels increased the challenge of speech perception we performed an ANOVA using the aov_ez function in the ‘afex’ package (Singmann, Bolker, Westfall, Aust, & Ben-Shachar, 2021) with factors of Training (pre vs. post) and Channels (clear, 16, 8, 4).

To quantify the relationship between acoustic degradation and speech perception performance we fit a logistic psychometric function to the word report accuracy data (separately for each participant, for and pre- and post-training tests) using the ‘quickpsy’ package (Linares & Lopez-Moliner, 2006). The parameters of the logistic function were estimated using direct maximisation of the likelihood with the following equation:

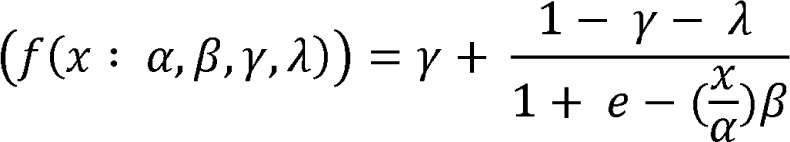

During the fitting, we treated clear speech as equivalent to 32-channel vocoded speech and converted the number of channels vocoded at each level of degradation into their log equivalents (χ). From each fit, we obtained alpha (α), the number of channels estimated to give 50% accuracy on the word report task. This value, referred to as “threshold number of channels” was used for the subsequent analyses of the impact of lesion on performance (c.f. McGettigan et al., 2014). Lower alpha values indicate that fewer channels were required to reach this threshold, and thus reflect better performance or more accurate perception. Beta (β) corresponds to the slope or steepness of the curve. Gamma (γ) is the guess rate, which was fixed to 0 for this open set speech task. Lambda (λ) is the lapse rate, or expected proportion of errors as the number of channels reaches the highest levels. Lambda represents the upper horizontal asymptote and was fixed at 1 minus the proportion of correct word report observed for clear speech for each participant separately.

### 3.5. Task 2. Semantically-ambiguous speech comprehension and adaptation

The second task increased speech comprehension difficulty at the lexical-semantic level, by the inclusion of semantically ambiguous words, in sentence contexts that in most cases supported the lower frequency meaning. We assessed speech comprehension in terms of the speed and accuracy of judging the coherence of these sentences, which were interspersed with sentences without ambiguities and anomalous sentences. The coherence judgement task appeared well-suited for assessing competence at semantic ambiguity resolution for several reasons. First, to respond accurately listeners must understand the whole sentence and not just identify one (or more) unusual words. For example, a sentence might initially make sense but then become anomalous only at the end (“*It was a rainy day and the family were thinking to the banana”)* or might initially seem odd but would eventually make sense *(“It was a terrible hand and the gambler was right to sit it out”)*. Secondly, since most of the meanings that we used in the sentences were the less frequent meanings, accurate performance relies on listeners utilising contextual cues to select the appropriate meaning rather than the higher-frequency, more accessible meaning. The use of lower-frequency word meanings also maximised our chance to observe word meaning priming effects, as described below. Thirdly, participants make a speeded judgement giving a continuous measure of performance in addition to accuracy.

To assess the increase in availability of low frequency word meanings in response to experience, we measured changes to meaning preferences in a word association task. This task provides a direct measure of how participants interpret ambiguous word forms in the absence of any sentence context. Specifically, using two counterbalanced sentence sets, we measured the increase in proportion of word association responses that were consistent with the (low frequency) meaning used in the sentence context for ambiguous words that had been heard (primed) compared to those that had not (unprimed). Counterbalanced assignment of sentences to primed and unprimed conditions for different participants ensured that differences in meaning frequency or dominance did not confound assessment of the word-meaning priming effect (for further discussion of word-meaning priming, see Rodd et al., 2013).

#### 3.5.1. Stimuli

The stimuli for the coherence judgement task were one hundred and twenty declarative sentences, selected from two previous studies [DETAILS REMOVED FOR DOUBLE BLIND REVIEWING]. Of these, 40 were high-ambiguity coherent sentences, 40—low-ambiguity coherent sentences and 40—anomalous sentences. The high-ambiguity sentences each contained 2 ambiguous words that were disambiguated within the sentence (e.g., “*The PITCH of the NOTE was extremely high*”; the ambiguous words were not repeated across the set of 40 sentences). Prior dominance ratings (Gilbert & Rodd, 2022) indicated that in most of the sentences, the context biased the interpretation of the ambiguous words towards their subordinate (less frequent) meanings (mean dominance = 0.31; SD = 0.25). The low-ambiguity sentences were matched with the high-ambiguity sentences across the set for number of words, number of syllables, syntactic structure and naturalness but contained words with minimal ambiguity (e.g. “*The pattern on the rug was quite complex*”). These 80 coherent sentences were separated into two lists (List A and List B), each containing 20 high-ambiguity and 20 low-ambiguity sentences. Participants were presented with sentences from either list (List A or List B) and thus were exposed to half of the ambiguous words in this part of the experiment. Each list also contained all 40 anomalous sentences (i.e. the same sentences were presented to all participants) which had been created from the low-ambiguity sentences by randomly substituting content words matched for syntactic class, frequency of occurrence and numbers of syllables (e.g., “*There were tweezers and novices in her listener heat*”). Thus, the anomalous sentences had identical phonological, lexical and syntactic properties but lacked coherent meaning (see Table 2 for psycholinguistic properties of the 3 sentence types).

**Table 2.**
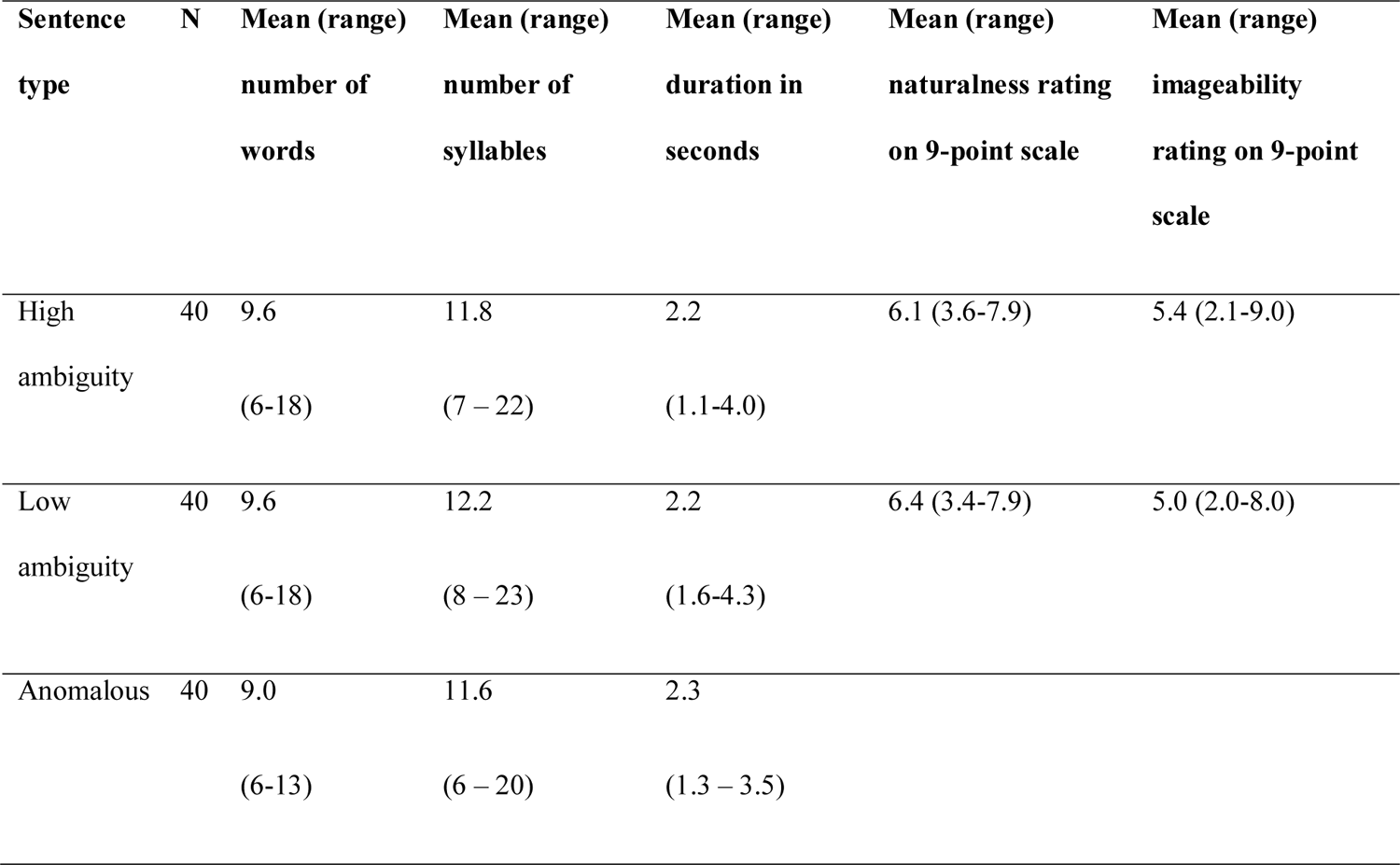
Descriptive characteristics of the three sentence types.

The stimuli for the word association task were the 80 ambiguous target words from the 40 high-ambiguity sentences. Given that participants had only heard half of the high-ambiguity sentences in the sentence coherence judgement task (List A or List B), for 40 of the ambiguous words, the subordinate meaning was primed (previously heard in a supportive sentence context) and for the other 40, the subordinate meaning was not primed.

Sentences and single words were recorded individually by a male native speaker of British English (MHD) and sentences were equated for RMS amplitude across conditions.

#### 3.5.2. Procedure

The task consisted of two phases. In the first phase, participants listened to 80 sentences (20 high-ambiguity, 20 low-ambiguity, 40 anomalous) and had to judge as quickly and as accurately as possible the coherence of each sentence. They indicated their response by pressing a green button if the sentence made sense and a red button if it did not. Participants were given examples (not included in the test set) to encourage them to listen to the sentence in its entirety before making the judgment.

Following the coherence judgement task, participants completed other behavioural tasks (not relevant to the current investigation) for 20-30 minutes before moving to the second phase: a word association task. In this phase, participants heard 80 ambiguous words presented in isolation, of which half had been presented in phase 1 (primed) and half were new (unprimed; counterbalanced across participants). For each word, participants had to repeat it and then say the first related word that came to mind. Responses were audio recorded and later coded as consistent with the subordinate meaning (e.g., “*NOTE*-music”) or inconsistent with the subordinate meaning (“*NOTE*-write”).

#### 3.5.3. Data processing and analysis

For each participant, we assessed whether they could discriminate the coherent sentences (high-ambiguity and low-ambiguity) from the incoherent sentences better than would be expected by chance, by calculating d-prime values for the high-ambiguity and low-ambiguity sentences separately:

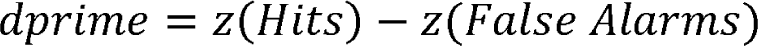

Hits correspond to the proportion of coherent sentences correctly judged as coherent and false alarms—to the proportion of incoherent sentences incorrectly judged as coherent. To allow for calculation of the z-scores, hit rates of 1 were adjusted by 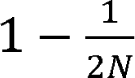 (i.e., to a value of 0.975) and false alarm rates of 0 were adjusted by 1/2N (Macmillan & Kaplan, 1985, i.e. to a value of 0.0125).

We analysed differences in accuracy and response times between the high-ambiguity and low-ambiguity sentence trials with a logistic mixed effects model (GLMM; accuracy) or a linear mixed effects model (LMM; log-10 response times). The models had a single categorical fixed effects predictor for Sentence Type (High-ambiguity or Low-ambiguity) with deviation coding defining one planned contrast: High-ambiguity = 1/2 versus Low-ambiguity = −1/2. The final models each contained a by-subject and by-item random intercept.

The correlational analyses used the model residuals (comparing predictions to the data) to estimate the ambiguity response time effect (difference between responses for high-ambiguity and low-ambiguity sentence trials) for each participant. A positive residual difference indicates that the participant’s ambiguity effect was larger than predicted by the model (response times were slower than estimated for the high-ambiguity condition and/or faster than estimated for the low-ambiguity condition). A negative residual difference means that their response time effect was smaller than predicted by the model (response times were faster than estimated for the high-ambiguity condition and/or slower than estimated for the low-ambiguity condition).

For the word association task, each response was independently coded for consistency with the subordinate meaning used in the priming sentence by two of the authors (LM and ZB), who were blind to the experimental condition (primed/unprimed) of the responses. For example, the word “ball” came from the sentence “*The ball was organised by the pupils to celebrate the end of term*”, so responses such as “party” and “dance” were coded as consistent whereas responses such as “kick” and “round” were coded as inconsistent. The consistency scores for the unprimed words give a baseline measure of the preference for the dominant meaning. Response codes from the first author were used with the exception of one participant for whom data were lost and only the codings from the second rater were available; inter-rater reliability for the remainder of the responses was high (94% agreement from 1,360 responses, Cohen’s Kappa = 0.862).

We analysed differences in the proportions of responses consistent with the subordinate meaning between primed and unprimed words (word meaning priming) with a logistic mixed effects model (GLMM) with a categorical fixed effect predictor for Priming Type (Primed or Unprimed) with deviation coding defining one planned contrast: Primed = 1/2 versus Unprimed = −1/2. There was also a continuous fixed effect predictor of Meaning Dominance (obtained from the prior dominance ratings (Gilbert, Betts, Jose, & Rodd, 2017)) and the associated interactions. The final model contained a by-subject and by-item random intercept and a by-subject random slope for Dominance.

In the main correlational analyses we used the model residuals (comparing predictions to the data) to estimate word priming effects (difference between response values for primed and unprimed words) for each participant. A positive residual difference indicates that the participant’s priming effect was larger than predicted by the model (proportion of responses consistent with the subordinate meaning was underestimated for the primed condition and/or overestimated for the unprimed condition). A negative residual difference means that their priming effect was smaller than predicted by the model (proportion of responses consistent with the subordinate meaning was overestimated for the primed condition and/or underestimated for the unprimed condition).

## 4. Results

### 4.1. Task 1. Acoustically degraded speech perception and adaptation

#### 4.1.1. Word Report Task

Figure 2A shows the mean proportion of words correctly reported for speech with different numbers of channels, for the pre- and post-training tests. Word report accuracy was near ceiling (100%) for the clear speech reflecting the participants’ ability to perform the task, and decreased as the number of channels decreased (*F*(1,3) = 211.865, *p* < .0001) reflecting the challenge of the acoustic degradation, reaching floor levels of near 0% for the 4-channel vocoded condition. Accuracy was greater following training (*F*(1,3) = 32.287, *p* < .0001) showing that participants were able to learn. The interaction between training and number of channels was not significant (*F*(1,3) = 2.358, *p* = .083).

**Figure 2.**
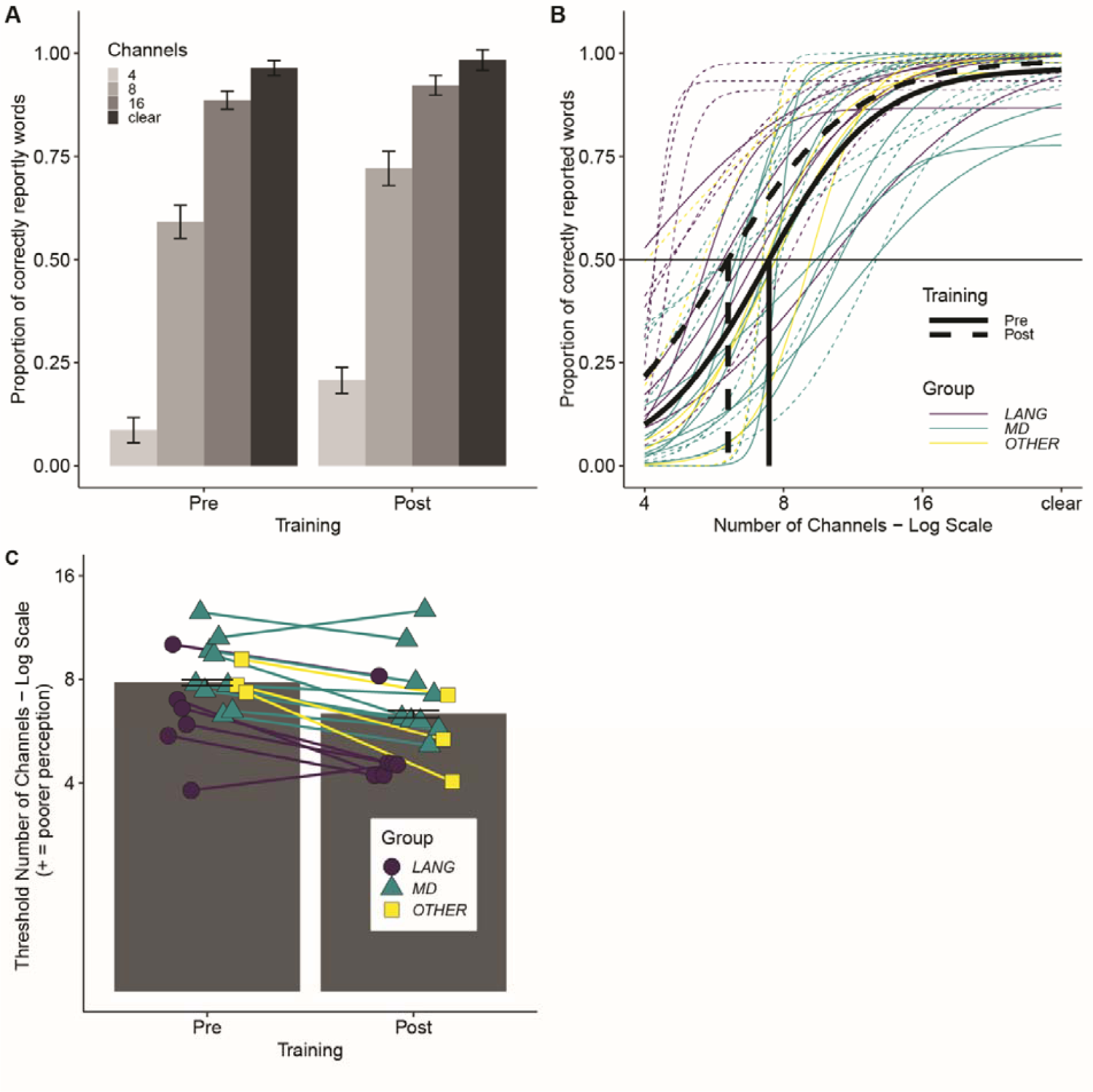
A. Word report accuracy scores for different levels of degradation and the Pre- and Post-Training tests separately. Bars show mean values across all 18 participants and error bars show ±1 SEM, adjusted to remove between-subject variance (Morey, 2008). B. Psychometric logistic function fits separately for the pre-(solid) and post-(dashed) training data for the mean across all 18 participants (black colour) and each participant separately (coloured by Group). The horizontal line indicates the 50% word report accuracy threshold. Vertical lines indicate the estimated threshold number of channels corresponding to the 50% word report accuracy threshold for the mean fits across all 18 participants. C. Estimated threshold number of channels (log scale) required for 50% accuracy in the word report task for the Pre- and Post-Training tests separately. Bars show mean values across all 18 participants and error bars show ±1 SEM, adjusted to remove between-subject variance (Morey, 2008). Individual participant values are overlaid (colour and shape reflect participant Group; see supplementary materials).

The outputs of fitting the data with a logistic psychometric function are shown in Figure 2B. Analyses to assess the impact of lesions on performance used the threshold number of channels (the estimated number of channels required for 50% word report accuracy) with lower values reflecting better perception (fewer channels needed to reach 50% accuracy). Figure 2C shows the mean performance before and after training for the group and for individual participants.

Figure 3 shows the relationship between degraded speech perception performance and the extent and location of lesions. Correlational analyses showed that the mean threshold number of channels across pre- and post-training tests positively correlated with damage to the MD network (*r* =.427, *p* = .039) but not with damage to the Language network (*r* = − 0.152, *p* = .727), or with total damage (*r* = 0.216, *p* = .194). Comparisons of these correlations demonstrated that poorer speech perception was numerically, but not significantly more strongly predicted by damage to the MD network than to the Language network (*z* = −1.954, *p* = .051). There was no evidence for MD network damage being more predictive of speech perception than total damage (*z* = −1.122, *p* = .262). There were no correlations between perceptual learning of degraded speech (i.e. change in threshold from pre- to post-training) and the volume of brain damage in the language network (*r* = −0.123, *p* = .727), MD network (*r* = 0.003, *p* = .504) or total damage (*r* = −0.006, *p* = .491. There was also no evidence that MD damage was more predictive of degraded speech perception than of degraded speech adaptation (*z* = −1.403, *p* = .161).

**Figure 3.**
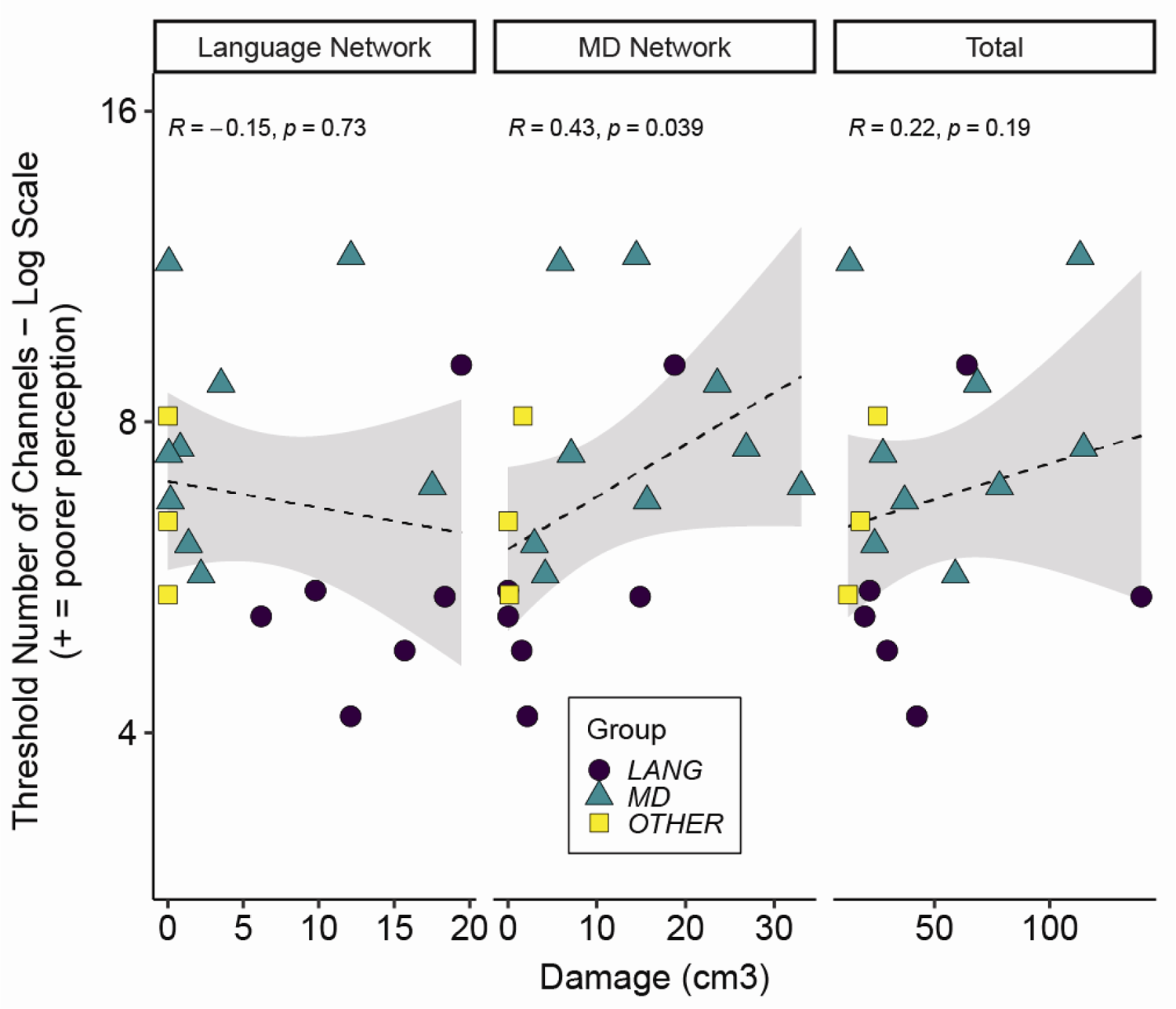
Individual participant data for estimated threshold number of channels (log scale) required for 50% word report accuracy for the mean of Pre- and Post-Training tests plotted separately against damage to the language network, to the MD network and total damage (colour and shape reflect participant Group; see supplementary material). Higher threshold number of channels indicates worse speech perception performance. The dashed line shows the linear best fit and grey shaded areas show 95% confidence intervals.

### 4.2. Task 2. Semantically ambiguous speech comprehension and adaptation

#### 4.2.1. Sentence Coherence Judgment Task

##### Accuracy Analysis

Of the 1,440 experimental trials (18 participants X 80 items) we excluded three trials with very fast responses (more than 300 ms before the offset of the sentence), which were assumed to arise from accidental key presses or anticipatory responses. This resulted in the exclusion of 2 anomalous sentence trials and 1 low-ambiguity sentence trial.

All participants showed d-prime values substantially above 0 indicating successful discrimination of the incoherent sentences from both the high-ambiguity (mean = 3.66, SD = 0.25, range = 3.0 – 4.20) and the low-ambiguity sentences (mean = 3.75, SD = 0.37, range = 2.72 – 4.20). Participant mean d-prime values across all sentences showed no correlation with lesion volume in the language or MD networks or with total lesion (*p*s > .7).

As the false alarm rate was necessarily identical for high-ambiguity and low-ambiguity conditions (since we only included a single set of incoherent sentences), differences in accuracy between the high-ambiguity and low-ambiguity sentences can be assessed using error rates when participants judged these coherent sentences to be anomalous. Therefore, for the main accuracy analyses we excluded the 40 anomalous sentence trials, leaving 719 trials (1 trial was excluded based on RTs; see above). Across all participants the proportions of correct responses were near ceiling (mean = 0.97, SD = 0.03, range = 0.92 – 1.0). Mean error rates can be seen for the different Sentence Types and for individual participants in Figure 4A. The mixed effect model showed no effect of Sentence Type (model coefficient: β = −0.440, SE = 0.538, *z* = −0.817, *p* = .414) and hence we have no evidence that sentences containing ambiguous words were less well understood.

**Figure 4.**
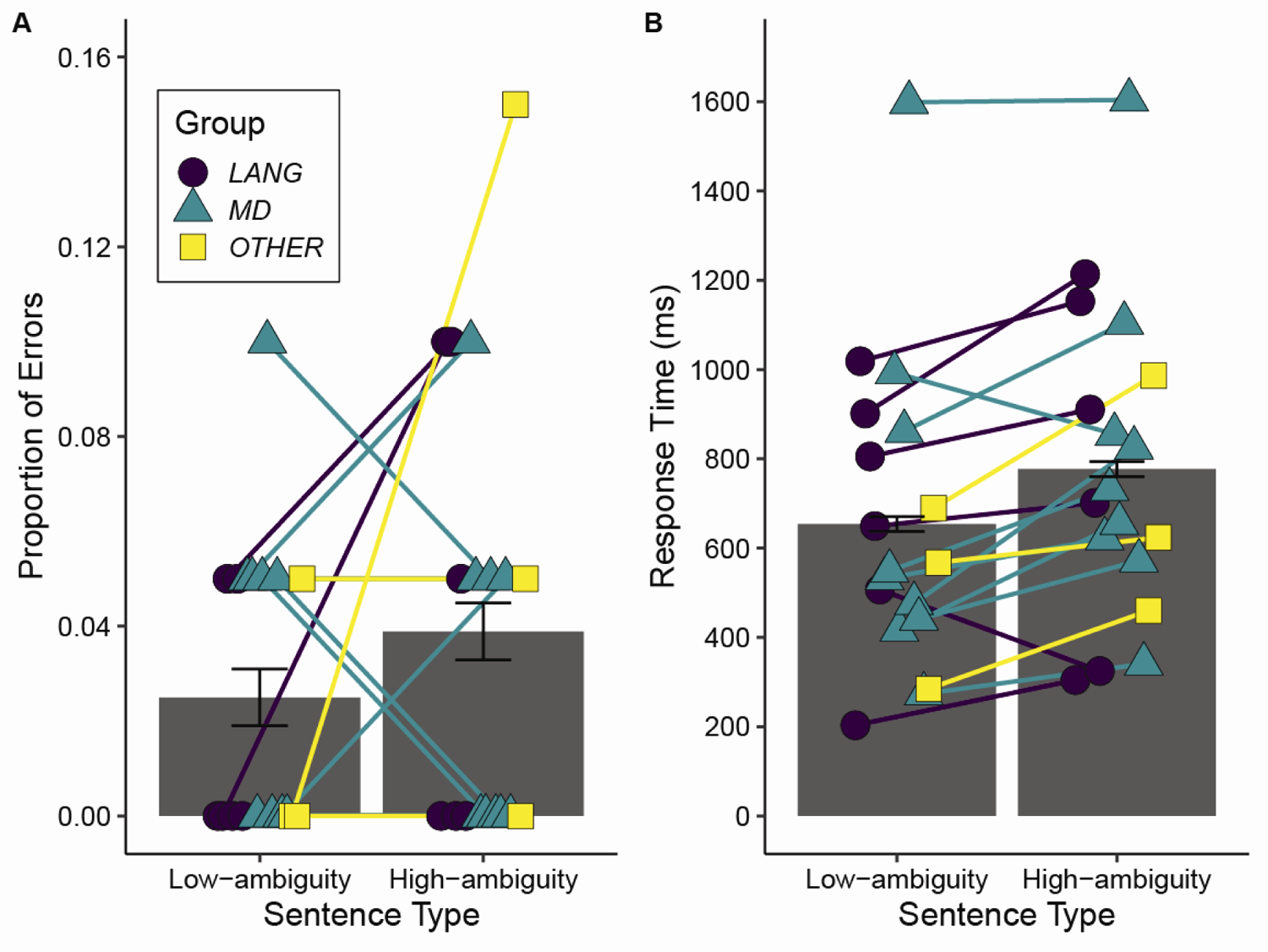
A. Proportion of errors and B. Response times measured from sentence offset for coherence judgements to low-ambiguity and high-ambiguity sentences. In each case, bars reflect the mean values across all 18 participants and error bars show ±1 SEM, adjusted to remove between-subject variance (Morey, 2008). Individual participant values are plotted (colour and shape reflect participant Group; see supplementary materials).

##### Response Time Analysis

The response time analyses focused on ambiguous and unambiguous sentence trials. Of the 720 total experimental trials (18 participants X 40 items), we excluded trials incorrectly judged as incoherent (23 trials: 14 ambiguous, 9 unambiguous). For exclusions of trials based on response times, we followed the general principle of minimal trimming with model criticism (Baayen & Milin, 2010). We excluded trials with very fast response times (less than 300 ms before offset; as for the accuracy analysis), which were assumed to reflect accidental key presses (1 trial) as well as trials with very slow response times (3 trials with responses longer than 4,000 ms after sentence offset) because we were interested in speeded responses. Further exclusions were considered after first determining whether any transformation of the dependent variable was required to meet assumptions of the LME models, of homogeneity of residual variance and normally distributed residuals. Model diagnostic plots (quantile-quantile and histogram plots of the residuals) for the raw, log10-transformed and inverse transformed response time data showed that log10 transformation best met the assumptions. Examination of the plots for outliers indicated that no further trimming was necessary, thus there were 693 correctly judged coherent trials included in the analyses. Figure 4B shows the mean response times for the different Sentence Types and participants.

The mixed effect model confirmed that high-ambiguity sentences were responded to more slowly than low-ambiguity sentences (β = 0.043, SE = 0.019, *t*(2.319) = 2.319, *p* = .023), showing that ambiguous words increased the challenge of sentence comprehension (Figure 4B). However, there was no correlation between individual ambiguity response time effects (model residual difference measure) and extent of damage to the language network (*r* = −0.102, *p* = .656), the MD network (*r* = −0.038, *p* = .559) or overall damage (*r* = 0.076, *p* = .383).

#### 4.2.2. Word Association Task

We excluded primed trials corresponding to words from sentences that were responded to incorrectly in the coherence judgement task. This resulted in exclusion of 28 trials (words from 14 ambiguous sentences: 1.94% of data) across the 18 participants, leaving 1,412 observations. For unprimed words (i.e., for ambiguous words that were not presented to participants in the coherence judgement task), the mean proportion of responses (across items and participants) that were consistent with the subordinate meaning of the word was 0.29 (SD = 0.09). This value, which gives a baseline measure of the preference for the dominant meaning, indicates that the sentence-primed meanings were indeed the subordinate or less preferred meanings (note that the value is similar to the one derived from an existing database (Gilbert & Rodd, 2022; see section on Stimuli above). Figure 5 shows the mean proportion of responses consistent with the subordinate meaning for primed and unprimed words.

**Figure 5.**
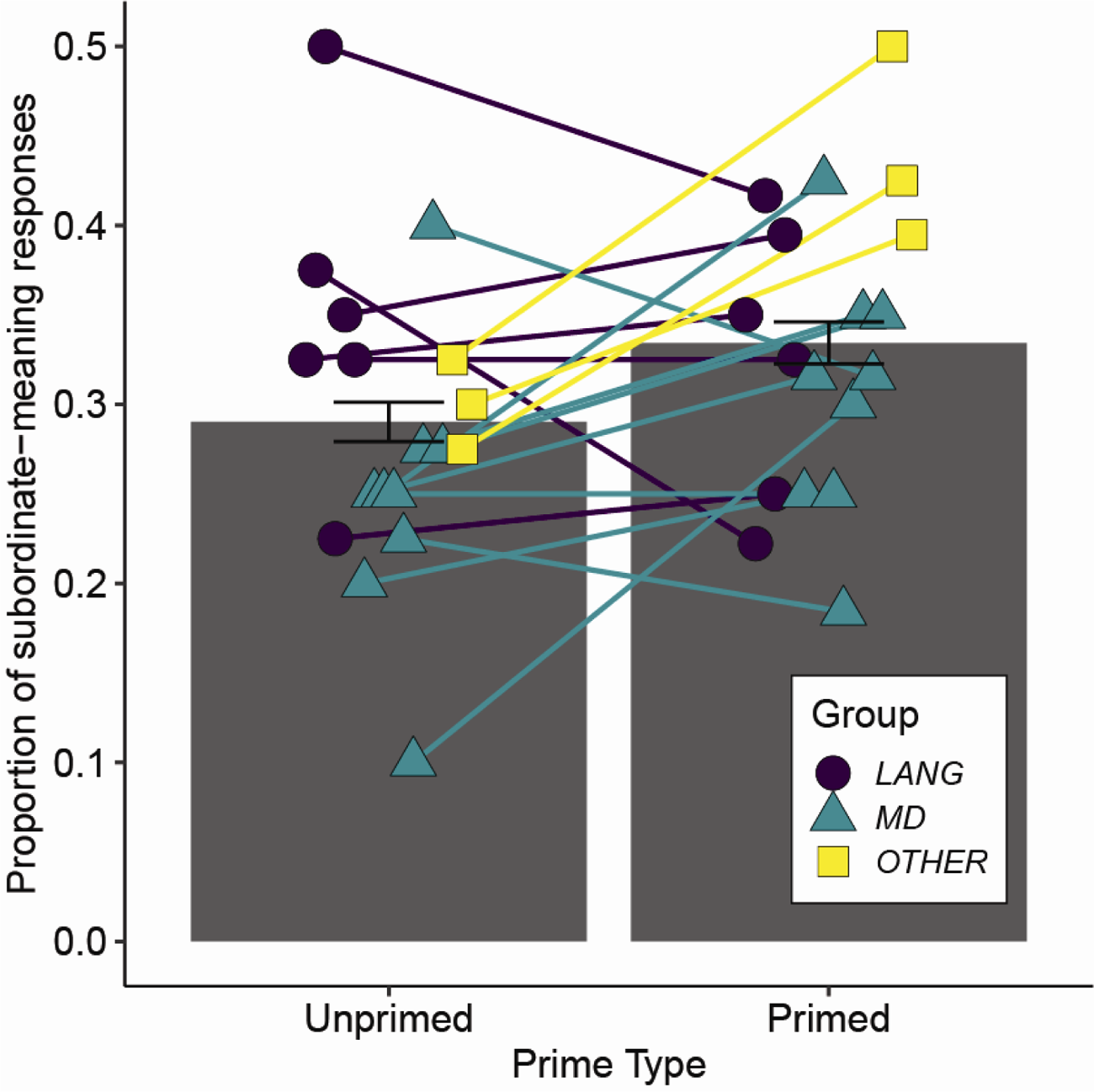
Proportion of responses consistent with the subordinate meaning for ambiguous words in the Unprimed and Primed conditions. Bars reflect the mean values across all 18 participants and error bars show ±SEM, adjusted to remove between-subject variance. Individual participant values are plotted (colour and shape reflect participant Group; see supplementary materials).

We observed a main effect of Priming (β = 0.352, SE = 0.137, *z* = 2.565, *p* = .010), which reflects a higher proportion of responses consistent with the subordinate meaning for the primed compared to unprimed words. This finding demonstrates a change in word meaning preferences in response to recent experience of sentences containing ambiguous words. We also observed a main effect of meaning Dominance of the word (β = 1.112, SE = 0.111, *z* = 10.054, *p* < .0001), reflecting an increase in proportion of responses consistent with the subordinate meaning as the dominance of that meaning increased (became less subordinate and closer in frequency to the alternative dominant meaning). There was an interaction between Dominance and Priming (β = −0.300, SE = 0.171, *z* = −2.124, *p* = .034), reflecting a stronger Priming effect for meanings that were more subordinate.

Correlational analyses revealed a negative relationship between individual word meaning priming effects (model residual difference measure) and the extent of damage to the language network (*r* = −0.659, *p* = .001) but not the MD network (*r* = −0.035, *p* = .446) or total damage (*r* = −0.180, *p* = .237). Comparisons between these correlations showed that word meaning priming was more strongly predicted by damage to the language network than to the MD network (*z* = −2.182, *p* = .0291) although the correlation between language network damage and word meaning priming was not significantly stronger than with total damage (*z* = −1.863, *p* = .062). There was also evidence that damage to the language network was more predictive of individual participants’ word meaning priming than the ambiguity response time effect (*z* = −2.6523, *p* = .008). Figure 6 shows scatter plots of the correlations between word-meaning priming and damage to language, MD Networks, and total damage. A further correlational analysis showed no relationship between participants’ ambiguity response time effect and their word meaning priming effect (both measured using the model residuals: *r*=-0.26, *p* = .298, 2-tailed), suggesting that reduced word meaning priming effect could not be simply explained as due to poorer comprehension.

**Figure 6.**
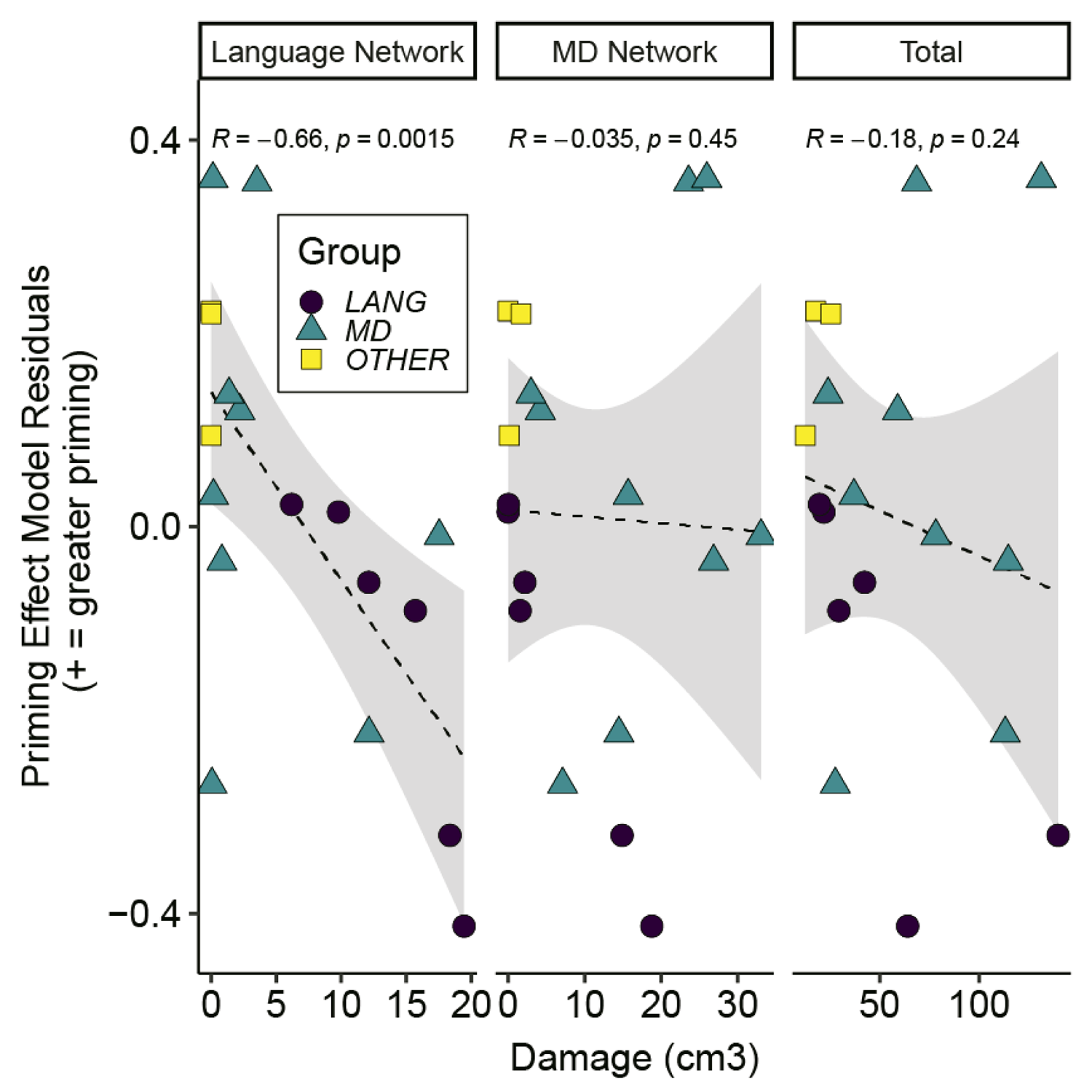
Individual participant data for word meaning priming effects estimated from the model residuals, plotted separately against damage to the language network, to the MD network and total damage (colour and shape reflect participant Group; see supplementary materials). The dashed line shows the linear best fit and grey shaded areas show 95% confidence intervals.

### 4.3. Dissociations between challenges to speech perception and comprehension

Tables 3A and 3B summarise the correlations between the lesion volume in each network and measures of perception, comprehension and adaptation, for the two types of challenge. Table 3A displays lesion-behaviour correlations from the 18 participants tested for each challenge and highlights comparisons between correlations with behaviour for lesions to the two networks and within each challenge type. Table 3B displays lesion-behaviour correlations for the 17 participants for whom we have data for both types of challenge. This table shows comparisons between correlations for the two types of challenge.

**Table 3.** Summary of correlations between the extent of damage to the two networks of interest and the tasks for the two types of challenge, and associated comparisons of the strength of these correlations. Significant lesion-behaviour correlations (in bold) are shown between the MD network and acoustically degraded speech perception (turquoise) and between the language network and semantic ambiguity adaptation (word meaning priming; purple). Arrows indicate significant differences (2-tailed) in the strength of pairs of correlations p<.05 (solid lines) and p<.1 (dashed lines). A. Correlations for data from the 18 participants for each type of challenge and associated within-challenge (across-lesion, across-task) comparisons. These are the results reported in the sections 4.1 and 4.2. B. Correlations for data from the 17 participants for whom we have data for both types of challenge and whose data contribute to the across-challenge comparisons. For all these results, negative correlations between lesion volume and task performance consistently indicates that an increase in lesion volume is associated with worse behavioural outcomes. Note that to achieve this consistency, we altered the sign of correlations for two of the tasks: (1) the perception of acoustically degraded speech since for the original analysis lower perceptual thresholds indicate better perception; (2) the comprehension of semantic ambiguity since previously a smaller response time difference between low and high ambiguity sentences indicates better comprehension.

|  |  | LESION |  |
| --- | --- | --- | --- |
| CHALLENGE |  | MD network | Language network |
| **Acoustic degradation**<br>$n=18^a$ | Perception | $r = -.427, p = .039$ | $r = .152, p = .727$ |
| | Adaptation | $r = .003, p = .504$ | $r = -.123, p = .313$ |
| **Semantic ambiguity**<br>$n=18^b$ | Comprehension | $r = .038, p = .559$ | $r = .102, p = .656$ |
| | Adaptation | $r = -.035, p = .446$ | $r = -.659, p = .001$ |

B
|  |  | LESION |  |
| --- | --- | --- | --- |
| CHALLENGE |  | MD network | Language network |
| **Acoustic degradation**<br>$n=17^{a,b}$ | Perception | $r = -.530, p = .014$ | $r = .058, p = .587$ |
| | Adaptation | $r = -.002, p = .500$ | $r = -.140, p = .297$ |
| **Semantic ambiguity**<br>$n=17^{a,b}$ | Comprehension | $r = -.064, p = .403$ | $r = .174, p = .748$ |
| | Adaptation | $r = -.201, p = .220$ | $r = -.639, p = .003$ |

<sup>a</sup> excluding participant md6<sup>b</sup> excluding participant md10— $p < .05$ , 2-tailed- - - $p < .10$ , 2-tailed

As detailed in the task-specific results (summarised in Table 3A), degraded speech perception was predicted by the degree of MD network damage and this correlation was in the opposite direction to, but not significantly stronger than, the correlation with language network damage volume (see section 4.1.1). Conversely word meaning priming was predicted by language network damage and this correlation was significantly stronger than the non-significant correlation with MD network damage volume (section 4.2.2). This double dissociation provides evidence for causal associations between the integrity of the MD network and abilities at degraded speech perception and between the language network and word meaning priming. Further comparisons of the strength of correlations between tasks within the same type of challenge (acoustic degradation or semantic ambiguity) showed that damage to the language network was more predictive of impaired word meaning priming than of comprehension of ambiguous sentences. This finding provides support for a specific contribution of the language network to adaptation that is independent to its role in comprehension (at least as measured here), which was shown to be largely independent of language or MD network lesions. However, there was no evidence that damage to the MD network was more strongly predictive of degraded speech perception than of degraded speech adaptation.

To further explore the specificity of the contribution of the MD and language networks to acoustically degraded speech perception and word meaning priming respectively, we also compared the strength of correlations between tasks using data from the 17 participants who performed both the acoustic degradation and the semantic ambiguity tasks (Table 3B). There was no evidence that damage to the MD Network was more predictive of degraded speech perception than of word meaning priming (*z* = −1.042, *p* = .300), or of comprehension of ambiguous sentences (*z* = 1.644, *p* = .100). However, damage to the language network was more predictive of impaired word meaning priming than of degraded speech perception (*z* = −2.160, *p* = .031), although it was not significantly more predictive of word meaning priming than of degraded speech adaptation (*z* = −1.678, *p* = .093).

## 5. Discussion

We report two main findings. First, we show that damage to the domain-general MD network, but not the language-selective network, causes significant impairments to the perception of acoustically degraded speech. Word report accuracy for noise-vocoded sentences decreased as the number of channels in the vocoder decreased reflecting an increased challenge to speech perception. The degree of perceptual impairment (i.e. the number of channels required for 50% correct word report) depends on the extent of damage to the MD network, but not damage to the language network (Figure 3; Table 3). Word recognition improved following a period of training, reflecting adaptation or perceptual learning for this form of acoustic degradation, but the degree of learning was not reliably predicted by lesion location or extent.

In contrast to these results with acoustically challenging speech, we found no evidence that semantically-challenging speech comprehension was dependent on the MD system: all participants were highly accurate in judging the coherence of sentences and were no less accurate when the sentences contained ambiguous words, indicating an intact ability to access the typically less frequent (subordinate) word meanings used in our high-ambiguity sentences. Although participants were slower to make judgements for sentences which include ambiguous words, reflecting more effortful comprehension when words have multiple meanings, there was no significant association between response time slowing for ambiguous sentences and the extent of damage to the MD or language networks.

Our second main finding is that despite accurate comprehension of semantically ambiguous speech, damage to the language, but not to the MD network caused a significant reduction in updating of word meaning preferences following recent linguistic experience. As shown in previous studies of individuals without brain lesions, our participants were (as a group) more likely to generate a word associate related to the less frequent meaning of an ambiguous word when they had encountered this meaning in an earlier sentence (word meaning priming, as reported by Rodd et al., 2013). However, the magnitude of this word meaning priming effect was predicted by the extent of damage to the language, but not the MD network (Figure 6; Table 3), a dissociation that was supported by a statistically significant difference between the strength of these two correlations. The reduction in word meaning priming was not explained by sentence comprehension difficulties as there was no correlation between the magnitude of word meaning priming and increased response times when judging the coherence of sentences containing semantically ambiguous words. Furthermore, across-task comparisons showed that the damage to the language network was more predictive of impaired word meaning priming than impaired comprehension of ambiguous sentences or impaired perception of acoustically degraded speech.

Below, we discuss our two main findings in greater detail. First, we discuss possible cognitive operations performed by the MD network that are required for the perception of acoustically degraded speech. We then turn to the linguistic challenge of resolving lexical-semantic ambiguity. We discuss the functional contribution of the language-selective network in adaptation such that low frequency meanings of semantically ambiguous words become more accessible following recent exposure. In a final section we consider the dissociation between these different challenges to speech processing and explore implications for the neural basis of speech perception, comprehension, and adaptation.

### 5.1. The MD network makes a causal contribution to perception of acoustically degraded speech

Recently it has been argued that the MD network does not play a functional role in language comprehension (Blank & Fedorenko, 2017; Diachek et al., 2020; Shain et al., 2020; Wehbe et al., 2021; for reviews, see Fedorenko, 2014; Campbell & Tyler, 2018) and that activations observed during language comprehension within the MD network reflect a generic increase in effortful processing or contributions to specific task demands (such as decision making or button presses). However, this line of research left open the possibility that MD contributions may be necessary when speech is acoustically degraded and challenging to perceive (Diachek et al., 2020). Here we provide novel evidence that the MD network indeed makes a causal contribution to perception of acoustically degraded speech.

Previous fMRI studies have shown that listening to acoustically challenging speech is associated with an increase in activation in prefrontal and motor regions that plausibly fall within the MD network (Adank, 2012; Davis & Johnsrude, 2003; Du et al., 2016; Erb et al., 2013; Hardy et al., 2018; Hervais-Adelman et al., 2012; Rysop et al., 2021; Vaden et al., 2015; Vaden et al., 2013; Wild et al., 2012). However, these studies did not explicitly define MD regions and hence this association has not been firmly established. A substantial advance, then, comes from our finding that neural integrity of the MD network supports more successful word report for degraded speech, which allows us to conclude a causal role of MD regions in degraded speech perception.

The MD network has previously been linked to a diverse range of domain-general cognitive constructs, including executive control, working memory, and fluid intelligence. These constructs may reflect a combination of different cognitive operations including setting and monitoring of task goals, directing attention, and the storage, maintenance, integration and inhibition of information across different time scales. It is therefore of interest to consider which of these operations, performed by the MD network, might be critical for the perception of acoustically degraded speech. For example, focused attention may be particularly important when the identities of specific phonemes or words are uncertain. Monitoring may be important for tracking the accuracy of phoneme perception and word recognition over time.

Future work can tease apart these possible distinct cognitive operations, either by focusing on potential contribution of distinct sub-networks within the broader MD network, or by exploring correlations between these other functions of MD networks and perception of degraded speech. Given strong evidence of inter-regional correlations during naturalistic listening paradigms (Assem et al., 2020; Blank et al., 2014; Mineroff et al., 2018; Paunov et al., 2019), we here treated the MD network as a functionally integrated system. However, other research concerned with domain general cognitive processes has proposed that the MD network consists of at least two interconnected, but distinct sub-networks (one comprising lateral frontal and parietal areas, and the other—cingular and opercular areas), which may contribute differently to cognition (Dosenbach, Fair, Cohen, Schlaggar, & Petersen, 2008; Dosenbach et al., 2007; Nomura et al., 2010). In the context of effortful speech comprehension, Peelle (2018) proposes a three-way distinction between fronto-parietal, premotor and cingular-opercular contributions to attention, working memory and performance monitoring processes respectively. Consistent with the proposed role of cingular-opercular regions are data showing that activation is associated with better word recognition on subsequent trials (Vaden et al., 2015; Vaden et al., 2013), which may reflect mechanisms for tracking the accuracy of phoneme perception and word recognition over time. Although the present data cannot adjudicate between bi- or tripartite views, further research using similar methods, and larger, or more carefully selected patient groups could potentially dissociate the effect of lesions of these three sub-networks and establish underlying mechanisms. For example, we might predict that focal damage to cingular-opercular regions would result in a greater impairment in degraded speech perception when perceptual difficulty varies from trial-to-trial compared to cases in which trial difficulty is grouped into blocks.

Replicating a range of previous behavioural findings (Davis et al., 2005; Hervais-Adelman et al., 2008; Huyck & Johnsrude, 2012; Loebach & Pisoni, 2008; Peelle & Wingfield, 2005; Sohoglu & Davis, 2016), we showed that listeners adapt to acoustically degraded speech over time. This finding extends earlier observations of perceptual learning to individuals with lesions to language-selective and domain-general regions. We found no evidence that damage to either MD or language-selective networks led to reduced perceptual learning, and hence cannot make causal claims about the contribution of either network to this form of learning. Nonetheless, this null finding does not rule out a causal role for region(s) within these networks that could be tested in future. It is possible that sub-components of the language or MD networks, or other auditory and sensorimotor brain regions, might be critical for perceptual learning. Previous fMRI research provides some support for each of these possibilities, with learning effects associated with increased activity in the IFG, precentral gyrus and cerebellum (Eisner, McGettigan, Faulkner, Rosen, & Scott, 2010; Guediche et al., 2014; Hervais-Adelman et al., 2012), while MEG has shown reduced post-learning responses in the Superior Temporal Gyrus (auditory speech cortex; Sohoglu & Davis, 2016). Future research could focus on each of these candidate regions and further test potential dissociations between their contributions to speech perception and learning.

Although we could not separate the significant predictive effect of MD damage on perception from the (null) effect of MD lesions on learning, previous research has explored neural and cognitive distinctions between the mechanisms driving the use of prior context in the recognition of acoustically degraded words and subsequent learning (Hardy et al., 2018; Sohoglu & Davis, 2016). This is also an exciting avenue for future exploration using similar methods as those employed here but with larger groups of participants with more varied lesions.

### 5.2. The language-selective network makes a causal contribution to adaptation to semantically ambiguous speech

Semantically ambiguous words introduce a substantial challenge to speech comprehension because of the need to engage competition processes to select between alternative meanings and the cognitive cost of reinterpretation when initial selection fails (Rodd et al., 2002; Rodd et al., 2010, 2012). The presence of two or more ambiguous words in each of the high ambiguity sentences used in our study made comprehension especially challenging. Nonetheless, we observed that comprehension – indicated by judging high ambiguity sentences to be coherent – was ultimately successful (although slower than for low ambiguity sentences) and that accuracy in judging coherence did not differ between high and low ambiguity sentences. Neither response time differences nor the accuracy of coherence judgements were associated with the degree of damage to MD or language-selective brain networks. Thus, our study does not provide evidence for a specific causal role of either of these brain networks for comprehension of sentences containing ambiguous words.

Despite intact comprehension of sentences containing semantically ambiguous words, we observed differential effects of lesion location and extent on learning mechanisms involved in adapting lexical-semantic processing after successful disambiguation. Previous research has established that recent exposure to low-frequency (subordinate) meanings of ambiguous words in a sentence context facilitates subsequent meaning access and selection of those meanings, a process termed word-meaning priming (Betts et al., 2018; Gaskell et al., 2019; Gilbert, Davis, Gaskell, & Rodd, 2018; Rodd et al., 2016; Rodd et al., 2013). Previous functional imaging studies have not studied neural activity associated with word-meaning priming and hence the present results make a novel contribution to understanding the neural basis of this adaptation process. We here replicated the standard word meaning priming effect for the group of participants tested overall, but showed that the magnitude of the priming effect was significantly reduced by damage to the language-selective but not the MD network.

There is substantial anatomical overlap between the language network shown here to be critical for updating of word-meaning preferences following successful disambiguation, and the fronto-temporal brain regions previously shown to respond to semantic ambiguity resolution (Bilenko et al., 2009; Musz & Thompson-Schill, 2017; Rodd et al., 2005; Vitello et al., 2014; Zempleni et al., 2007; for a review, see Rodd, 2020), consistent with shared neural resources between semantic comprehension and subsequent adaptation. The absence of an effect in the present study of language-network damage on immediate comprehension, coupled with the observed dissociation between the contribution of the language-network to semantic adaptation and immediate comprehension may therefore reflect the relatively high functioning of the volunteers, the limited severity of the language lesions, and/or the relative insensitivity of the comprehension task to distinguishing between these relatively unimpaired volunteers. It also remains possible that particular subregions of the language network are differentially important for immediate comprehension compared to subsequent adaptation and this could be explored in future work.

One striking illustration of the longevity of learning is that word-meaning priming has previously been observed 24 hours after a single exposure to an ambiguous word; especially if there is an intervening period of sleep (Gaskell et al., 2019). This latter finding, in combination with a wider literature on the role of consolidation processes that facilitate the acquisition of new lexical knowledge (Dumay & Gaskell, 2007; Gaskell & Dumay, 2003; Tamminen, Payne, Stickgold, Wamsley, & Gaskell, 2010), led Gaskell et al. (2019) to suggest that word meaning priming may involve a two-stage complementary systems account of learning (McClelland, 2013), as proposed for the acquisition of novel words (Davis & Gaskell, 2009). According to this account, short-term learning arises from hippocampally-mediated binding of associations between words in the sentences, while these short-term changes are consolidated into long-term changes to word meaning preferences after sleep.

The present study constrains these complementary systems accounts of learning by revealing a causal contribution of language-selective cortical regions even for short-term adaptation of familiar word meanings. Future work could further consider the interaction of hippocampal and cortical regions in the learning and maintenance of meaning preferences over different time scales and the relationship between learning novel vocabulary and updating of existing lexical semantic knowledge (for recent meta-analyses of word form learning and consolidation, see, Schimke, Angwin, Cheng, & Copland, 2021; Tagarelli, Shattuck, Turkeltaub, & Ullman, 2019). We note that previous research has shown that individuals with aphasia (identified behaviourally) can learn novel vocabulary but that learning is highly variable (Kelly et al., 2009; Tuomiranta et al. 2011; 2014). The present work similarly shows variability in the impact of cortical lesions on adapting the meanings of familiar words. However, our participants were not recruited on the basis of language impairment and retained good comprehension both on a standardised measure of sentence comprehension (TROG2) and on the experimental measure of ambiguity resolution tested here. It might be that individuals with more extensive lesions to language selective cortex, or more focal lesions of posterior temporal and inferior frontal regions that contribute to ambiguity resolution would show a greater impairment to comprehension. Such a finding would suggest that a common set of cortical regions support comprehension and learning of ambiguous words in sentences. Further refinement of lesion definitions and tests of larger samples of individuals could also provide more detailed anatomical evidence concerning the relative contributions of language-selective and/or domain-general sub-regions of the IFG which lie in close proximity (Fedorenko & Blank, 2020) and that have not been dissociated in previous imaging research on semantic ambiguity resolution.

### 5.3. Neural dissociation of different challenges to speech perception and comprehension

Taken together, our findings provide a double dissociation indicating independent functional contributions of the MD and language-selective networks to responding to and adapting to different types of difficult-to-understand sentences. Specifically, we show that the challenge of perceiving acoustically degraded sentences (measured in terms of word report accuracy) is causally linked to the degree of damage to the MD network but not to the language-selective network (although the comparison of correlations was not statistically significant; see Table 3). Conversely, the challenge of post-comprehension adaptation to semantically ambiguous words in sentences (measured in terms of word meaning priming) causally depends on the integrity of the language-selective but not the MD network; moreover, in this case there is a reliable difference between the significant (language) and null (MD) correlations.

Here we tested a limited set of challenges to speech comprehension, thus we cannot make general statements concerning dissociable contributions made by each of these cortical networks to all forms of perceptual or semantic challenge. However, our data provide initial evidence for the task-specificity of causal contributions. Focusing first on the effect of language network damage, the correlation between language lesion volume and word meaning priming was significantly different from the null correlation between lesion volume and coherence judgment response times for ambiguous sentences, indicating a distinction between the contribution of the network to initial comprehension of and later adaptation to lexico-semantic ambiguity. As discussed above (Section 5.2) we are not suggesting that the language network is not important for comprehension, only that adaptation processes are perhaps more sensitive than comprehension to the integrity of the language network.

Furthermore, the reliable correlation between lesion volume and word-meaning priming could also be dissociated from the (null) effect of lesions on degraded speech perception, suggesting that the integrity of the language network may be more important for lexico-semantic than for acoustic challenges, at least for the group of patients tested here. However, the dissociation with the null effect of language network damage on degraded speech adaptation was not reliable.

Equivalent across-task comparisons of the effect of MD network damage did not reach statistical significance. Despite a reliable correlation between MD lesion volume and impaired perception of acoustically degraded speech, this effect could not be clearly dissociated from the null effects of lesion volume on tasks involving semantic processing, or perceptual and semantic adaptation. We therefore cannot draw strong conclusions about the specific contribution that the MD network makes to degraded speech perception based on the current data.

Further studies exploring a wider range of challenges to speech comprehension and with larger samples of participants might specify the causal contributions identified here in more detail. For example, we might assess lesion correlates of perception and adaptation for other forms of perceptual challenges to speech comprehension – e.g. using speech in background noise, or perceptually ambiguous speech sounds (see Mattys et al., 2012 for a review of these listening challenges). Future studies might also consider whether other forms of semantic, syntactic or lexical challenge to comprehension are also causally associated with the integrity of the MD or language networks. In this way, building on the current methods and findings, one could map the hierarchy of cognitive processes involved in speech perception and comprehension onto specific brain regions that support them. However, larger samples of patients will be needed if we are to conduct more anatomically specific analyses at the level of individual voxels or functional parcels rather than the larger networks studied here.

### 5.4. Conclusions

Speech comprehension in naturalistic situations requires listeners to accommodate and learn in response to a range of perceptual and semantic challenges that make spoken sentences more difficult to recognise and understand. Behavioural data from individuals with lesions to language-selective and domain-general MD networks demonstrate different functional contributions of these two networks depending on the source of the listening challenge. In particular, the MD network appears to be necessary for the perception of acoustically degraded speech, whereas using recent experience to update meaning preferences for ambiguous words appears to depend on anatomically distinct, fronto-temporal regions argued to form a specialised language network.

In this work we considered two specific challenges, but future work should consider whether differences in the ways in which acoustic degradation and lexical-semantic ambiguity engage and depend on the domain-general MD network and domain-selective language network translate to other perceptual and semantic challenges and to more naturalistic speech processing. For example, speech perception must be resilient in the face of unfamiliar accents, mispronunciations, and competing sounds. Comprehension processes must accommodate multiple forms of syntactically or semantically complex and ambiguous speech. Many of these are situations in which activation of inferior frontal regions has been observed (Blanco-Elorrieta, Gwilliams, Marantz, & Pylkkanen, 2021; Boudewyn et al., 2015; January, Trueswell, & Thompson-Schill, 2009; Kuperberg et al., 2003; Novais-Santos et al., 2007) and an attribution to domain-general MD processing has sometimes been made.

However, given the evidence for functionally distinct language-selective and domain-general sub-regions lying in close proximity within the IFG, and the individual variability in their precise locations (Fedorenko & Blank, 2020), such conclusions may be premature. Further studies of individuals with focal lesions can be used to determine whether accommodating these other perceptual and semantic challenges to speech processing similarly depends on the integrity of domain-general or language-selective brain regions. These perceptual and semantic challenges are common for the noisy and ambiguous spoken language that listeners perceive and comprehend every day.

## Acknowledgements

We thank Rahel Schumacher and Vitor Zimmerer for discussion and Peter Watson for statistical advice

## Conflict of interest

Authors report no conflict of interest

## Funding Sources

This work was supported by Medical Research Council Grants MC_UU_00005/5 to MHD and MC_UU_0005/6 to JD; by NIH Awards R01-DC016950 and R01-DC016607 to EF; by research funds from the McGovern Institute for Brain Research to EF. JR is funded by the Economic and Social Research Council Grant ES/S009752/1

## Notes

### Competing Interest Statement

The authors have declared no competing interest.

