## Supplementary Information for "Causal contributions of the domain-general (Multiple Demand) and the language-selective brain networks to perceptual and semantic challenges in speech comprehension"

**Supplementary materials**

### 1. Methods

#### 1.1. Participant group assignment

As detailed in the manuscript (section 3.2) we calculated the lesion volume falling into each network for each of the 19 participants. Participants were assigned to one of three groups (LANG, MD or OTHER) based on the proportion of the lesion falling into Language and Multiple Demand regions as well as the overall proportion of each network that was damaged (Figure S1). A participant was assigned to either the “LANG Group” or the “MD Group” depending on whether a greater proportion of their lesion was in the Language or the MD network, so long as damage extended to at least 1% of the given network. If less than 1% of the most affected network was damaged, the patient was assigned to the “OTHER” Group. A participant was also assigned to the “OTHER” Group in the case that less than 1% of both networks were affected and less than 1% of the lesion fell in each network.

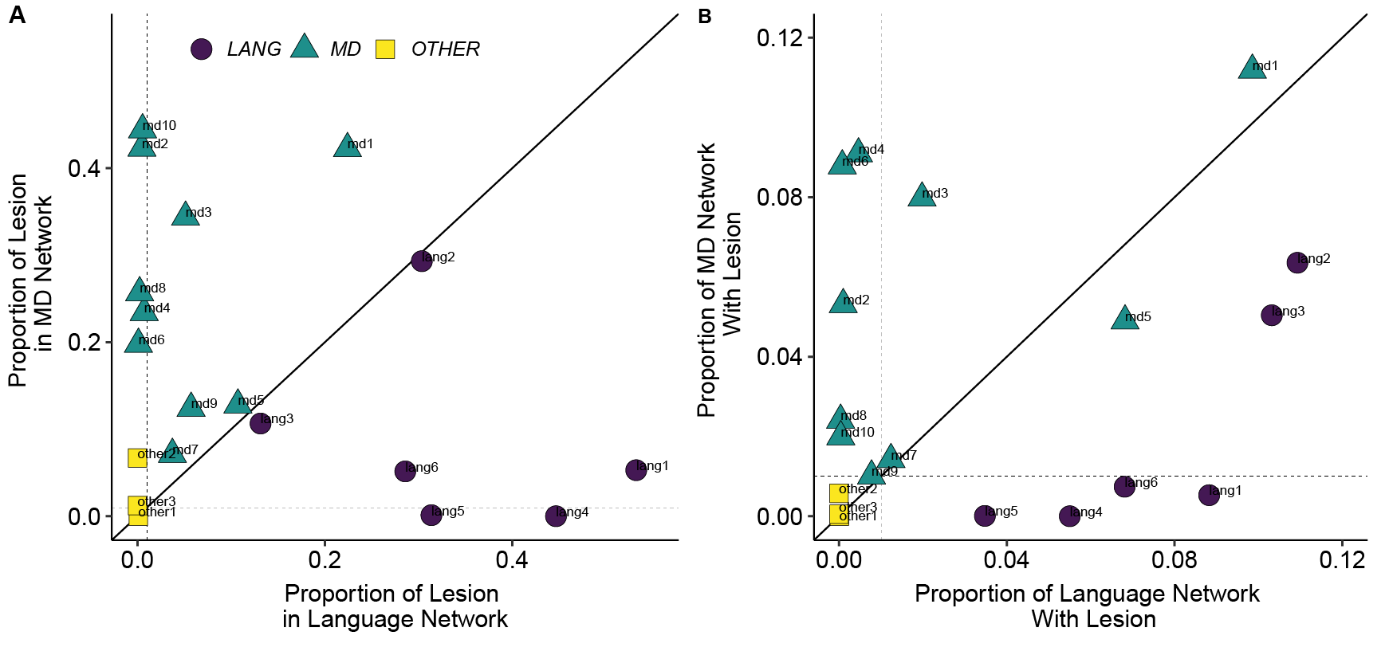

Figure S 1 Proportion of lesion falling into each network for each of the 19 participants in the present study. Solid line depicts an equal proportion of lesion falling into each network and shows the categorisation of the participants into the LANGUAGE (LANG) and MD Groups. Dotted lines show 1% threshold, which was used for categorisation of participants into the OTHER Group.

#### 1.2. Statistical Analyses

We assessed differences in behavioural responses across the different experimental conditions and patient groups by fitting either logistic or linear mixed effects models (GLMM or LMM) using the ‘lme4’ package (Bates, Machler, Bolker, & Walker, 2015). As the linear mixed effects model summaries produced by the ‘lme4’ package do not output significance values for fixed effects, significance of the individual model coefficients was determined with Satterthwaite’s approximation for degrees of freedom using the ‘lmerTest’ package (Kuznetsova, Brockhoff, & Christensen, 2015) because this method gives acceptable Type I error rates with these models (Luke, 2017).

In addition to the fixed effect predictors for our experimental conditions of interest, we used the maximal random effects structure that could be supported by the data because inclusion of all random slopes for fixed effects helps reduce the likelihood of Type I errors/ false positives (Barr, Levy, Scheepers, & Tily, 2013). If the maximal model did not converge, we removed the by-item and by-subject random slopes then intercepts. If these models did not converge, we removed the random effects term that accounted for the least variance, continuing in this way until the model converged. All tests of fixed effects were evaluated using models with this random effects structure. We evaluated the significance of main effects and interactions in two ways: (1) we performed model comparisons using χ^2^ values from likelihood ratio tests to compare the full fixed effect model to a model without the interaction term or without the main effect term; (2) we obtained the significance of the individual model coefficients from the *z*-test (GLMM) or *t*-test (LMM) statistics in the model summary. Significant interactions were followed up with pairwise comparisons using χ^2^ values with Holm p-value adjustment for multiple comparisons implemented in the ‘phia’ package (Rosario-Martinez, 2015).

### 2. Group Results

#### 2.1. Task 1. Acoustically degraded speech perception and adaptation

##### 2.1.1. Word Report Task

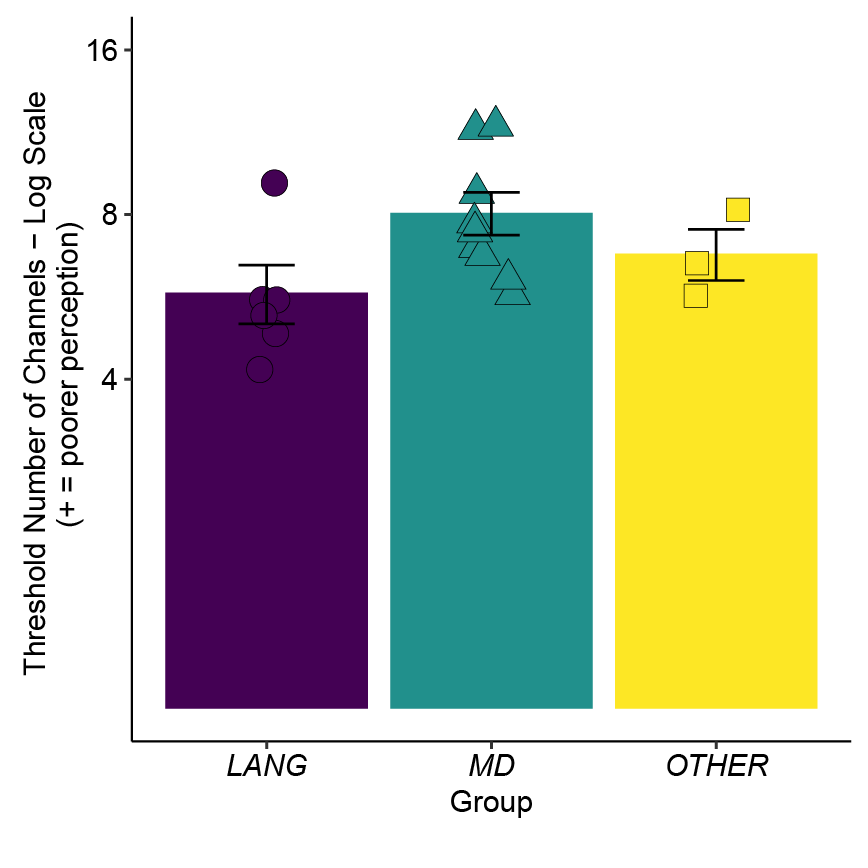

Figure S 2 Estimated threshold number of channels (log scale) required for 50% accuracy in the word report task for the mean of Pre- and Post-Training tests, for the participants in the LANG, MD, and OTHER Groups separately. Bars show mean values across the participants in each Group and error bars show ±1 SEM, adjusted to remove between-subject variance (Morey, 2008). Individual participant values are overlaid (colour and shape reflect participant Group). In each case, fewer channels indicate higher word report accuracy and thus better speech perception.

We analysed the estimated number of channels required for 50% word report accuracy to assess differences between the pre- and post-training word report and between participant groups with a logistic mixed effects (LME) model, which included categorical fixed effect predictors for Training (Pre or Post) and Group (Language, MD or Other) and the associated interaction. Within Training, deviation coding was used to define the one planned contrast: Post = 1/2 versus Pre = -1/2 and within Group, Helmert coding was used to define two planned contrasts: (a) Language =-1/3 versus MD = -1/3 versus Other = 2/3 and (b) Language = 1/2 versus MD = -1/2. The final model contained a by-subject random intercept.

The threshold number of channels required for 50% word report accuracy differed between the three groups (Figure S2, model comparisons: *χ*^2^(4) = 10.541, *p*  = .032). Specifically, it was greater for the MD compared to Language Group (model coefficients: *β* = -0.497, SE = 0.174, *t* = -2.850, *p* = .011) reflecting poorer speech perception (mean across pre- and post-training tests) for the MD Group. No reliable difference was observed between the Language and MD vs. Other Group (model coefficients: *β* = 0.004, SE = 0.21, *t*(18) = 0.02, *p* = .984). There was also a main effect of training such that threshold number of channels was lower for the post- compared to the pre-training test (model coefficient: *β* = 0.384, SE = 0.07, *t*(18) = 5.893, *p* < .001; model comparisons: *χ*^2^(3) = 19.451, *p*  < .001). Post-hoc tests indicated perceptual learning in each of the three groups (Language: *χ*^2^(1) = 11.363, p = .001, MD: *χ*^2^(1) = 7.828, p = .005, Other: *χ*^2^(1) = 15.847, p <.001). There was no evidence that the learning differed between the three groups overall, as indicated by the absence of a statistical interaction between Group and Training (model comparisons: *χ*^2^(2) = 3.806, *p*  = .150), or that learning differed between Language and MD (model coefficients: *β* = 0.286, SE = 0.159, *t*(18) = 1.802, *p* = .088) or between Language and MD vs. Other (model coefficients: *β* = 0.111, SE = 0.132, *t*(18) = 0.842, *p* = .411).

#### 2.2. Task 2. Semantically ambiguous speech comprehension and adaptation

##### 2.2.1. Sentence Coherence Judgment Task

For the coherence judgement task we analysed accuracy and response times to assess the differences between the high-ambiguity and low-ambiguity sentence trials and between the three participant groups. Differences were assessed with a logistic mixed effects model (GLMM; accuracy) or a linear mixed effects model (LMM; log-10 response times), with categorical fixed effects predictors for Sentence Type (High-ambiguity or Low-ambiguity) and Group (Language, MD or Other) and the associated interaction. Within Sentence Type, deviation coding defined one planned contrast: High-ambiguity = 1/2 versus Low-ambiguity = -1/2. Within Group, Helmert coding was used to define two planned contrasts: (a) Language =-1/3 versus MD = -1/3 versus Other = 2/3 and (b) Language = 1/2 versus MD = -1/2. The final models contained a by-subject and by-item random intercept.

**Accuracy Analysis.** There was no evidence for a main effect of Sentence Type (model comparisons: χ^2^(3) = 3.09, *p* = .386, model coefficient: *β* = -0.732, SE = 0.609, z = -1.202, *p* = .229), nor for a main effect of Group (model comparisons: *χ^2^*(4) = 2.49, *p* = .646, model coefficients: Language and MD vs. Other Group: *β* = -0.062, SE = 0.717, *z* = -0.086, *p* = .932; Lang vs. MD Group: *β* = 0.111, SE = 0.583, *z* = 0.191, *p* = .848). The Sentence Type X Group interaction was also not significant (model comparisons: *χ*^2^(2) = 2.368, *p* = .306, model coefficients: Sentence Type X Lang and MD vs. Other: *β* = -1.088, SE = 1.303, *z* = -0.836, *p* = .403, Sentence Type X Lang vs. MD: *β* = -1.178, SE = 1.068, *z* = -1.103, *p* = .270). Thus we have no evidence that sentences containing ambiguous words were less well understood, or that comprehension success differed between the three participant Groups.

**Response Time Analysis.** We observed a main effect of Sentence Type indicating slower responses to ambiguous sentences (model coefficients: *β* = 0.047, SE = 0.019, *t*(88.8) = 2.429, *p* = .017, model comparisons: *χ*^2^(3) = 6.341, *p* = .096). There was no main effect of Group (model comparisons: *χ*^2^(4) = 1.729, *p* = .786, model coefficients: Language and MD vs. Other: *β* = -0.053, SE = 0.072, *t*(18.43) = -0.738, *p* = .470; Language vs. MD: *β* = -0.007, SE = 0.060, *t*(17.92) = -0.124, *p* = .902). The Sentence Type X Group interaction was not significant (model comparisons: *χ*^2^(2) = 1.523, *p* = .562, model coefficients: Sentence Type X Lang and MD vs. Other: *β* = 0.030, SE = 0.031, *t*(667) = 0.966, *p* = .334; Sentence Type X Lang vs. MD: *β* = -0.013, SE = 0.023, *t*(601) = -0.572, *p* = .568). Thus, as expected, ambiguous words increased the challenge of sentence comprehension, but there was no evidence that the different participant Groups were differentially challenged.

##### 2.2.2. Word Association Task

First, as expected, we observed a main effect of meaning Dominance of the word (model coefficients: *β* = 1.217, SE = 0.123, *z* = 9.936, *p* < .0001, model comparisons: χ^2^(6) = 85.526, *p*  < .0001), reflecting higher proportion of responses consistent with the subordinate meaning as the dominance of this meaning increased (became less subordinate). Also as expected, was a main effect of Priming (model coefficient: *β* = 0.435, SE = 0.16, *z* = 2.778, *p* = .005; model comparisons: *χ*^2^(6) = 14.378, *p*  = .002), reflecting a higher proportion of responses consistent with the subordinate meaning for the primed compared to unprimed words (word meaning priming effect). There was also some evidence for an interaction between Dominance and Priming (model coefficients: *β* = -0.300, SE = 0.171, *z* = -1.754, *p* = .080, model comparisons: χ^2^(5) = 14.055, *p*  = .015), reflecting greater word meaning priming (relative increase in meaning preference for the subordinate meaning) for less dominant meanings.

There was a significant interaction between Priming and Group (Figure S3; model comparisons: χ^2^(6) = 14.104, *p* = .023). Pairwise comparisons with Holm adjustment showed significant priming effects in the Other Group (χ^2^(1) = 5.238, *p*  = .044) and MD Group (χ^2^(1) = 5.959, *p*  = .044), but no evidence for a priming effect in the Language Group (χ^2^(1) = 0.045, *p*  = .832). However, there was no evidence in the model for a specific difference in the extent of word meaning priming for the Language vs. MD Group contrast (model coefficient: *β* = -0.552, SE = 0.30, *z* = -1.821, *p* = .069), nor for the Language and MD vs. Other Groups contrast (model coefficient: *β* = 0.623, SE = 0.41, *z* = 1.521, *p* = .123).

We also observed a main effect of Group (model comparison: χ^2^(8) = 20.467, *p*  = .009), specifically a difference between the Language and MD Groups (model coefficient: *β* = 0.461, SE = 0.19, *z* = 2.393, *p* = .012), reflecting group differences in overall preference for the subordinate meaning, but not between the Language and MD vs. Other Group (model coefficient: *β* = 0.302, SE = 0.24, *z* = 1.260, *p* = .208). There was no higher order interaction between Dominance, Priming and Group (model comparisons: χ^2^(4) = 9.33, *p*  = .053).

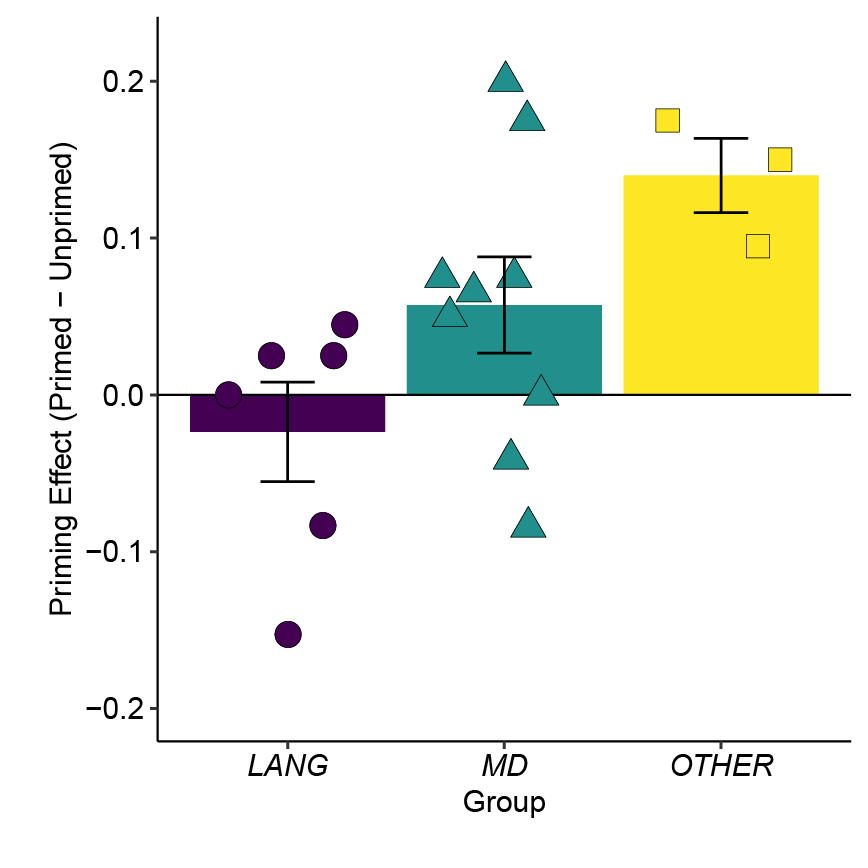

Figure S 3 Priming effect (proportion of responses consistent with the sentence-primed subordinate meaning for ambiguous words in the Primed minus the Unprimed conditions), for the 3 participant Groups separately. Bars reflect the mean values across the participants in each Group and error bars show ±1 SEM, adjusted to remove between-subject variance (Morey, 2008). Individual participant values are plotted separately (colour and shape reflect participant Group).

Barr, D. J., Levy, R., Scheepers, C., & Tily, H. J. (2013). Random effects structure for confirmatory hypothesis testing: Keep it maximal. *J Mem Lang, 68*(3).

Bates, D., Machler, M., Bolker, B. M., & Walker, S. C. (2015). Fitting Linear Mixed-Effects Models Using lme4. *Journal of Statistical Software, 67*(1), 1-48.

Kuznetsova, A., Brockhoff, P. B., & Christensen, R. H. B. (2015). Package "lmerTest".

Luke, S. G. (2017). Evaluating significance in linear mixed-effects models in R. *Behavior Research Methods, 49*, 1494-1502.

Morey, R. D. (2008). Confidence intervals from normalized data: A correction to Cousineau (2005). *Tutorial in Quantitative Methods for Psychology, 4*(2), 61-64.

Rosario-Martinez, H. D. (2015). phia: Post-Hoc Interaction Analysis (Version 0.2.1).
